# Reproductive phasiRNAs in grasses are compositionally distinct from other classes of small RNAs

**DOI:** 10.1101/242727

**Authors:** Parth Patel, Sandra Mathioni, Atul Kakrana, Hagit Shatkay, Blake C. Meyers

## Abstract

**Summary and keywords:**

- Little is known about the characteristics and function of reproductive phased, secondary, small interfering RNAs (phasiRNAs) in the Poaceae, despite the availability of significant genomic resources, experimental data, and a growing number of computational tools. We utilized machine-learning methods to identify sequence-based and structural features that distinguish phasiRNAs in rice and maize from other small RNAs (sRNAs).
- We developed Random Forest classifiers that can distinguish reproductive phasiRNAs from other sRNAs in complex sets of sequencing data, utilizing sequence-based (k-mers) and features describing position-specific sequence biases.
- The classification performance attained is >80% in accuracy, sensitivity, specificity, and positive predicted value. Feature selection identified important features in both ends of phasiRNAs. We demonstrated that phasiRNAs have strand specificity and position-specific nucleotide biases potentially influencing AGO sorting; we also predicted targets to infer functions of phasiRNAs, and computationally-assessed their sequence characteristics relative to other sRNAs.
- Our results demonstrate that machine-learning methods effectively identify phasiRNAs despite the lack of characteristic features typically present in precursor loci of other small RNAs, such as sequence conservation or structural motifs. The 5’-end features we identified provide insights into AGO-phasiRNA interactions; we describe a hypothetical model of competition for AGO loading between phasiRNAs of different nucleotide compositions.

## Introduction

Molecular and genomic studies coupled with deep sequencing have identified roles of many endogenous non-coding RNAs (ncRNAs) and small RNAs (sRNAs) at numerous developmental stages in many organisms (Tisseur *et al.*, 2011; Guttman & Rinn, 2012; Axtell, 2013; Kung *et al.*, 2013; Borges & Martienssen, 2015). Flowering plants have three major classes of sRNAs, all derived from ncRNAs: microRNAs (miRNAs), heterochromatic or Pol IV-dependent small interfering RNAs (P4-siRNAs), and phased, secondary, small interfering RNAs (phasiRNAs). This latter class has grown considerably with the discovery of germline-enriched, reproductive phasiRNAs most well described in the Poaceae, namely maize and rice (Johnson *et al.*, 2009; Komiya *et al.*, 2014; Zhai *et al.*, 2015b). Two classes of reproductive phasiRNAs are known: 21- nt pre-meiotic phasiRNAs that peak in abundance during somatic cell specification in maize (one week after anther initiation), and 24-nt meiotic phasiRNAs that peak during meiosis and are detectable until pollen maturation (one to two weeks after pre-meiotic phasiRNAs peak) (Zhai *et al.*, 2015b). The timing, localization, and narrow developmental time window of accumulation of the 21- and 24-nt phasiRNAs is conserved in rice and maize (Fei *et al.*, 2016). While the biogenesis and spatiotemporal patterns of accumulation of these reproductive phasiRNAs are now well described, our understanding of their function is still limited.

An analogy can be drawn between phasiRNAs of grass anthers and the PIWI-interacting RNAs (piRNAs) of animals, in aspects such as their biogenesis, developmental timing, and enrichment in reproductive organs. piRNAs play crucial roles in transposable element (TE) silencing and germline development from flies to fish to mammals (Meister, 2013). Yet, plants have a highly elaborate RNA-directed DNA methylation pathway (RdDM) that effectively silences most TEs (Matzke & Mosher, 2014), thus their need for yet another TE-silencing pathway is debatable. Emerging evidence implicates plant reproductive phasiRNAs in development; for example, MEL1, a rice Argonaute (AGO), is required for normal anther development (Nonomura *et al.*, 2007), and this AGO binds to 21-nt reproductive phasiRNAs (Komiya *et al.*, 2014). The functions and targets are yet to be determined for both 21- and 24-nt reproductive phasiRNAs, and it is not known whether they function in *cis* or *trans* (Song *et al.*, 2012a; Zhai *et al.*, 2015b). In fact, it is possible that they are merely decay products of more functionally relevant long ncRNA precursors. Understanding the role of phasiRNAs requires more detailed molecular and computational analyses that could also serve to direct future experiments. For example, identifying characteristic features or motifs that differentiate reproductive phasiRNAs from other sRNAs (miRNAs, P4-siRNAs, etc.) may provide clues as to their AGO loading or targets.

Work on animal piRNAs has used sequence-based characteristics to demonstrate their unique properties; significant insights have resulted from so-called *alignment-free approaches*. These methods use short nucleotide sequences, k-mers, and other features to distinguish between different types of sRNA sequences, and classify them into distinct groups. For example, Zhang *et al.*, 2011 developed a classifier that can distinguish piRNAs from non-piRNAs (miRNAs, snoRNAs, tRNAs, and lncRNAs) with precision over 90% and a recall over 60%, within a five-fold cross-validation. This work utilized data from five species including mice, humans, rats, fruit flies, and nematodes, effectively discriminating piRNAs. In a test of the validity of their classifier, Zhang *et al.* (2011) detected >87,000 of ∼130,000 piRNAs, in a total set of >600,000 sRNAs. Brayet *et al.* (2014) used a similar approach to identify piRNAs from sequences of several types (miRNAs, tRNAs, and 25-33 nt sequences from protein coding genes) in human and fruit flies with precision over 85% and a recall over 88%. As such, these alignment-free approaches are quite promising for characterizing subsets of sRNAs within large and complex pools of un-sorted sequences.

Our aim was to start with a set of known reproductive phasiRNAs (21- or 24-nt), develop and optimize a classification pipeline, and ultimately use this to sort previously unknown sequences from plants to find reproductive phasiRNAs from among other types of small RNAs. An additional product of this work was the sequence-based characteristics that comprise the output of the classifier, as these might identify novel aspects of reproductive phasiRNAs. In this work, we implemented machine-learning approaches to examine plant 21-nt pre-meiotic and 24-nt meiotic reproductive phasiRNAs, and to build a classifier that can automatically distinguish them from other sRNAs (i.e., miRNAs and P4-siRNAs). Our results provide insights into phasiRNA sequence composition profiles and biases, sequence-based and positional features, aspects of their biogenesis, features that may influence AGO sorting, predicted targets and possible functions.

## Methods

### Classification via machine learning

We use the Random Forest (“RF”) (Breiman, 2001) classification method, which is based on building an ensemble of decision trees. This method has proven effective for addressing a variety of classification problems in bioinformatics (Yang *et al.*, 2010; Lertampaiporn *et al.*, 2014). We employed the WEKA implementation of RF (Frank *et al.*, 2016) to build the model for distinguishing phasiRNAs (to which we refer as the *positive* set) from non-phasiRNAs (the *negative* set). As we study two sets of reproductive phasiRNAs, characterized by two distinct lengths, namely 21- and 24-nt, for each set we have trained two distinct classifiers, one for each length. When training each of these classifiers, we have varied the composition of the negative sets of non-phasiRNAs to which the phasiRNAs were compared (more details are in the data set used for cross validation study, Method S1).

To train and test the classifiers we developed, we have used the commonly used stratified five-fold cross-validation (CV) framework (Kohavi, 1995). Under this framework, the dataset is partitioned into five subsets, where each subset has the same ratio of positive instances to negative instances as the whole dataset. Once the data is partitioned, five iterations of training and testing are performed, where in each iteration four parts of the data (80%) are used for training and the remaining part (20%) is used for testing. To ensure stability and reproducibility of the results, the whole five-fold CV experiment was repeated five times, each using a different five-way split (partition) of the dataset.

### Performance evaluation

To assess classification performance we use the standard measures of *accuracy* (ACC), *specificity* (SP), *sensitivity* (SE), *positive predictive value* (PPV), and *area under the receiver operating characteristic curve* (AUC), whose formulae and descriptions are as follows:

- Sensitivity 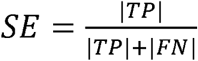;
- Specificity 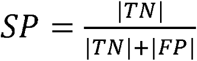;
- Accuracy ACC 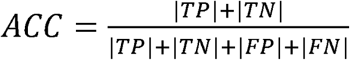;
- Positive Predictive Value PPV 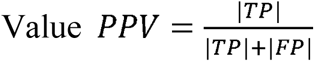; where *True Positives* (TP) denotes the set of correctly classified phasiRNAs, *True Negatives* (TN) denotes the set of correctly classified non-phasiRNAs, *False Positives* (FP) denotes the set of non-phasiRNA sequences that were classified as phasiRNAs, and *False Negatives* (FN) denotes the set of phasiRNA sequences that were not classified as such by our classifier. The number of items in the sets TP, TN, FP, and FN is denoted by |TP|, |TN|, |FP|, and |FN|, respectively.
- The Area Under the ROC Curve (AUC) is an effective and joint measure of sensitivity and specificity, which is calculated by the Receiver Operating Characteristic curve (ROC). AUC determines the relative performance of classifiers for correctly classifying phasiRNAs and non-phasiRNAs. Values of AUC are between 0 (worst performance) and 1 (best performance). ROC illustrates the true positive rate (sensitivity) against the false-positive rate (1 - specificity).

### Development of a machine learning classifier for plant small RNAs

The classification pipeline we developed takes as input a set of plant small RNA sequences to assess for each sequence whether it has attributes or not of a reproductive phasiRNA, based on a training/test set, returning a “yes” or “no” response. Thus, for this decision, feature characterization is crucial. The pipeline used several sequence- and structural-based features. One known feature of reproductive phasiRNAs is a 5’-terminal cytosine, described for 21-nt phasiRNAs bound by MEL1, a rice Argonaute (Komiya *et al.*, 2014). Another known characteristic of both 21- and 24-nt reproductive phasiRNAs is their origin from unique or low copy regions in the genome (Johnson *et al.*, 2009; Zhai *et al.*, 2015b). Beyond these features, little was known about their sequence composition, true even for other classes of plant small RNAs.

Thus, to build a classifier, we utilized an alignment-free approach based on k-mers. These k-mer motifs (more details in Method S2), together with the GC content and Shannon entropy of the small RNA, comprised the sequence-based features of the classifier. The other major component of the classifier was a set of positional features, calculated for each sequence to determine the presence or absence of a given nucleotide in a determined sequence position. These two sets of attributes for each sequence comprised 1498 features, most of which were short k-mers or words that we could use to classify plant small RNAs. Before each classification, feature selection of the top 250 most informative features (of the 1498) was performed as a step to better understand which features play key roles in classifying phasiRNAs; this allowed us to reduce the feature dimensionality comprising classification without compromising or negatively impacting the classifier’s performance (more details in Method S2). We have also experimented with different number of trees and of features used. Consequently, to estimate the performance of the classifier, RF was applied using 100 trees, five out of 250 features assessed (five randomly sampled features selected as candidates at each split) at each split, and five complete runs of the 5-fold CV.

The scripts used for this work are available on GitHub (https://github.com/pupatel/phasiRNAClassifier).

## Results

### Cross validation results distinguishes reproductive phasiRNAs from other sRNAs

We sought to identify unique attributes of rice and maize reproductive phasiRNAs relative to other, better-described small RNA classes. To do this, we developed a machine learning-based workflow focused on sequence-based and structural features of plant small RNAs (Fig. 1). To train the classifier, we used as positive examples known reproductive phasiRNAs from rice and maize, including both 21-nt and 24-nt phasiRNAs, while the negative sets consisted of P4-siRNAs, miRNAs, tRNAs, and rRNAs (see Method S1). We built and evaluated classifiers by utilizing different negative sets; the performance measurements were achieved via five-fold cross-validation (CV), and this 5-fold CV was completed five times on our datasets (see Methods for a more complete explanation). As noted above, the classification results were in terms of ACC, SE, SP, PPV, and AUC.

**Fig. 1.**
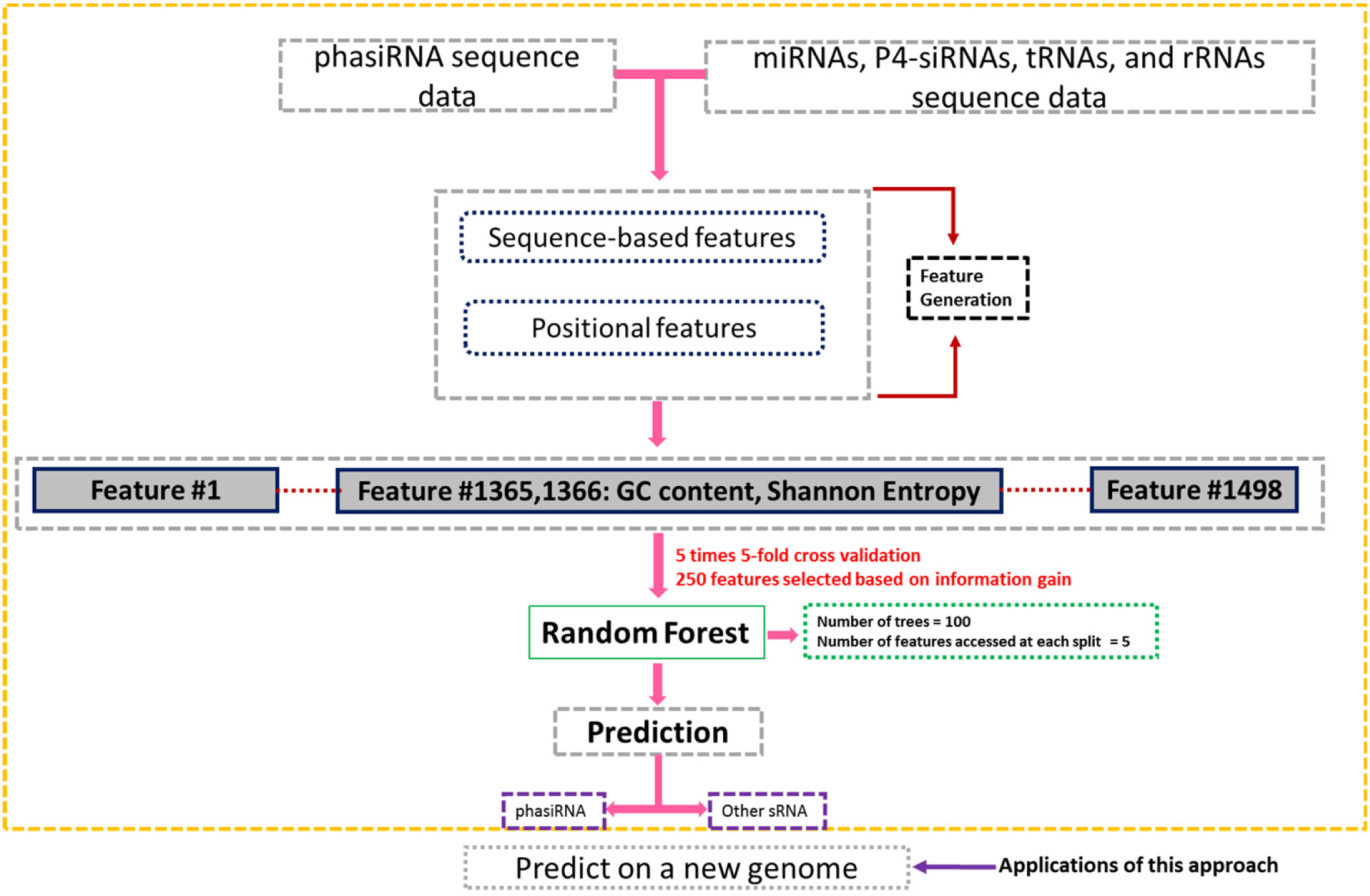
General workflow of our pipeline.

The results obtained from our classification pipeline using different negative sets, are shown in Table 1. The results, according to all performance measures, exceed 0.8 (with one exception, see below), for both 21- and 24-nt phasiRNAs. We first examined 21-nt phasiRNAs, and we compared phasiRNAs to a mixture of sRNAs that include selected miRNAs, P4-siRNA, tRNAs, and rRNAs; these latter four cases represent the four major negative sets (i.e. not phasiRNAs) found in a typical plant sRNA dataset (Table 1). In an initial comparison, the negative sets included miRNAs, tRNAs, and rRNAs of different lengths (randomly selected endogenous sequences); all P4-siRNAs were 24-nt. The classifier identified the combined negative set as quite distinct relative to 21-nt phasiRNAs (Table 1). In addition, we computed the area under the ROC curve (AUC) (Fig. S1c), demonstrating the performance of the above-mentioned classifier with an averaged AUC of 0.97. Next, we combined untrimmed miRNAs and 24-nt P4-siRNAs and still achieved high classification performance (Table 1). This classification result could indicate that length is a primary factor in classification, and thus we used trimmed negative sets to assess this possibility.

**Table 1.** Results of classification to distinguishing phasiRNAs of lengths 21-nt (top) and 24-nt (bottom) from other small RNA types.

| Classification |  | Performance Evaluation Measure |  |  |  |  |
| --- | --- | --- | --- | --- | --- | --- |
| Positive set | Negative set | ACC (±SD) | SP (±SD) | SE (±SD) | PPV (±SD) | AUC (±SD) |
| 21-nt phasiRNA | miRNAs* + P4-siRNA + tRNA* + rRNA* | 0.93 (±0.01) | 0.91 (±0.00) | 0.92 (± 0.01) | 0.93 (± 0.01) | 0.97 (± 0.00) |
| | miRNAs* + P4-siRNA | 0.93 (±0.01) | 0.90 (±0.00) | 0.94 (±0.01) | 0.87 (±0.02) | 0.97 (± 0.00) |
| | P4-siRNAs, 3' trimmed | 0.83 (±0.02) | 0.84 (±0.01) | 0.83 (±0.02) | 0.83 (±0.01) | 0.92 (± 0.00) |
| | miRNAs, 3' trimmed | 0.81 (±0.01) | 0.77 (±0.01) | 0.85 (± 0.03) | 0.78 (± 0.02) | 0.90 (± 0.01) |
| 24-nt phasiRNA | P4-siRNA | 0.84 (±0.01) | 0.82 (±0.00) | 0.84 (± 0.01) | 0.83 (±0.01) | 0.91 (± 0.01) |
| | miRNAs* + P4-siRNA + tRNA* + rRNA* | 0.87 (±0.01) | 0.82 (±0.00) | 0.91 (± 0.01) | 0.84 (±0.01) | 0.93 (± 0.01) |

Note: ACC, accuracy; SP, specificity; SE, sensitivity; PPV, positive predictive value. See methods for further detail. An asterisk (\*) next to a negative subset indicates no size selection or trimming of the sequences. Results are averaged over the five-fold cross-validation. The size of positive and negative sets are as follows: 21-nt phasiRNAs (n=2000), 24-nt phasiRNAs (n=2000), miRNA (n=756), P4-siRNAs (n=2000), tRNAs (n=500), and rRNAs (n=500).

We classified 21-nt phasiRNAs relative to 24-nt P4-siRNAs with 3 nt trimmed from the 3’ end (see Method S1); the classifier performed reasonably well (>0.8 for all measurements, ACC, SP, SE, PPV, and AUC). We also trimmed P4-siRNAs 3 nt from the 5’ end or from the internal 11^th^, 12^th^, and 13^th^ positions, observing no substantial changes in classification. We concluded that 21-nt reproductive phasiRNAs are compositionally distinct from P4-siRNAs. Finally, we related 21-nt phasiRNAs to 21-nt miRNAs (some trimmed, see Method S1), and found similar ACC, higher SE, but slightly lower SP and PPV; the lower SP may be attributed to fewer miRNAs (756 vs 2000 21-nt phasiRNAs in the positive set). This imbalance possibly misclassified some miRNAs, hence low specificity and a high number of false positives (lower PPV). We followed the same procedure in classifying the 24-nt phasiRNAs, first with 24-nt P4-siRNAs and next with the combined negative set. In both cases, the classification of the negative set against 24-nt phasiRNAs, resulted in strong scores for all four performance measurements (Table 1), again indicative that the 24-nt phasiRNAs are also compositionally distinctive. In addition, we observed an averaged AUC of 0.93 when classifying 24-nt phasiRNAs with the combined negative set (Fig. S1d). We concluded that our classification pipeline successfully classified reproductive phasiRNAs relative to other endogenous plant sRNAs with high values for ACC, SE, SP, PPV, and AUC.

Next, we investigated the predictive sensitivity of our pipeline, asking whether it can correctly classify previously unutilized members of a larger positive set of reproductive phasiRNAs. In other words, these new sequences were different from the 2000 used in the positive set during cross validation study. The classifier was given, first, 27500 21-nt phasiRNA sequences and, next, 7750 24-nt phasiRNA sequences (rice and maize combined, in each case). The classification pipeline based on models that combined each of the negative sets (miRNAs + P4-siRNAs + tRNAs + rRNAs) predicted 26208 21-nt phasiRNAs (SE > 0.96) and 7093 24-nt phasiRNAs (SE > 0.90), achieving high sensitivity in the two genomes from which we developed the models (Table 2a).

**Table 2.** Predictive performance of classification models of 21-and 24-nt phasiRNAs.

| a. Predictive sensitivity on rice and maize |  |  |  |
| --- | --- | --- | --- |
| Predictive sensitivity | Classification Model<br>(positive set vs negative set) | Performance Evaluation Measure |  |
|  |  | TP | SE |
| 21-nt phasiRNAs | 21-nt phasiRNA vs. miRNA + P4-siRNA + tRNA + rRNA | 26458/27500 | 0.962 |
| 24-nt phasiRNAs | 24-nt phasiRNA vs. miRNA + P4-siRNA + tRNA + rRNA | 7017/7750 | 0.905 |
| b. Cross-species predictive sensitivity in *Setaria viridis* |  |  |  |
| 21-nt phasiRNAs | 21-nt phasiRNA vs. miRNA + P4-siRNA + tRNA + rRNA | 1868/2000 | 0.934 |
| 24-nt phasiRNAs | 24-nt phasiRNA vs. miRNA + P4-siRNA + tRNA + rRNA | 1723/2000 | 0.861 |

Note: TP, true positive prediction; SE, sensitivity.

As additional test, we aimed to test the trained model in a different genome. To do so, we generated new small RNA data from panicles of the model grass *Setaria viridis* (see Method S3 and Table S1). We then applied the aforementioned classification models developed from rice and maize to assess reproductive phasiRNAs in these *S. viridis* data, to evaluate the potential of this approach across species. In *S. viridis,* a dataset and genome that we had not previously analyzed, the models predicted 1868 21-nt phasiRNAs and 1723 24-nt phasiRNAs with a sensitivity (SE) of > 0.93 and > 0.86, respectively (Table 2b). We concluded that the machine-learning method is effective for *de novo* classification of plant small RNAs.

### Position-specific biases in phasiRNAs relative to other small RNAs

Next, knowing that reproductive phasiRNAs are distinct from other classes of small RNAs, we sought to characterize these differences in greater detail, at the single nucleotide level. We computed single-nucleotide sequence profiles for the most abundant 1000 reproductive phasiRNAs (for 21-nt or 24-nt, rice and maize data combined), miRNAs, and 24-nt P4-siRNAs, determining the frequencies of each nucleotide (A, C, G, and U) at each position (Fig. 2). We then compared the position-specific base usage between the reproductive phasiRNAs and either miRNAs or 24-P4-sRNAs by conducting a two-tailed, rank sum test (*P* = 1e^−5^) to identify positions with statistically significant base usage that would distinguish phasiRNAs from either miRNAs or P4-siRNAs (Fig. 2).

**Fig. 2.**
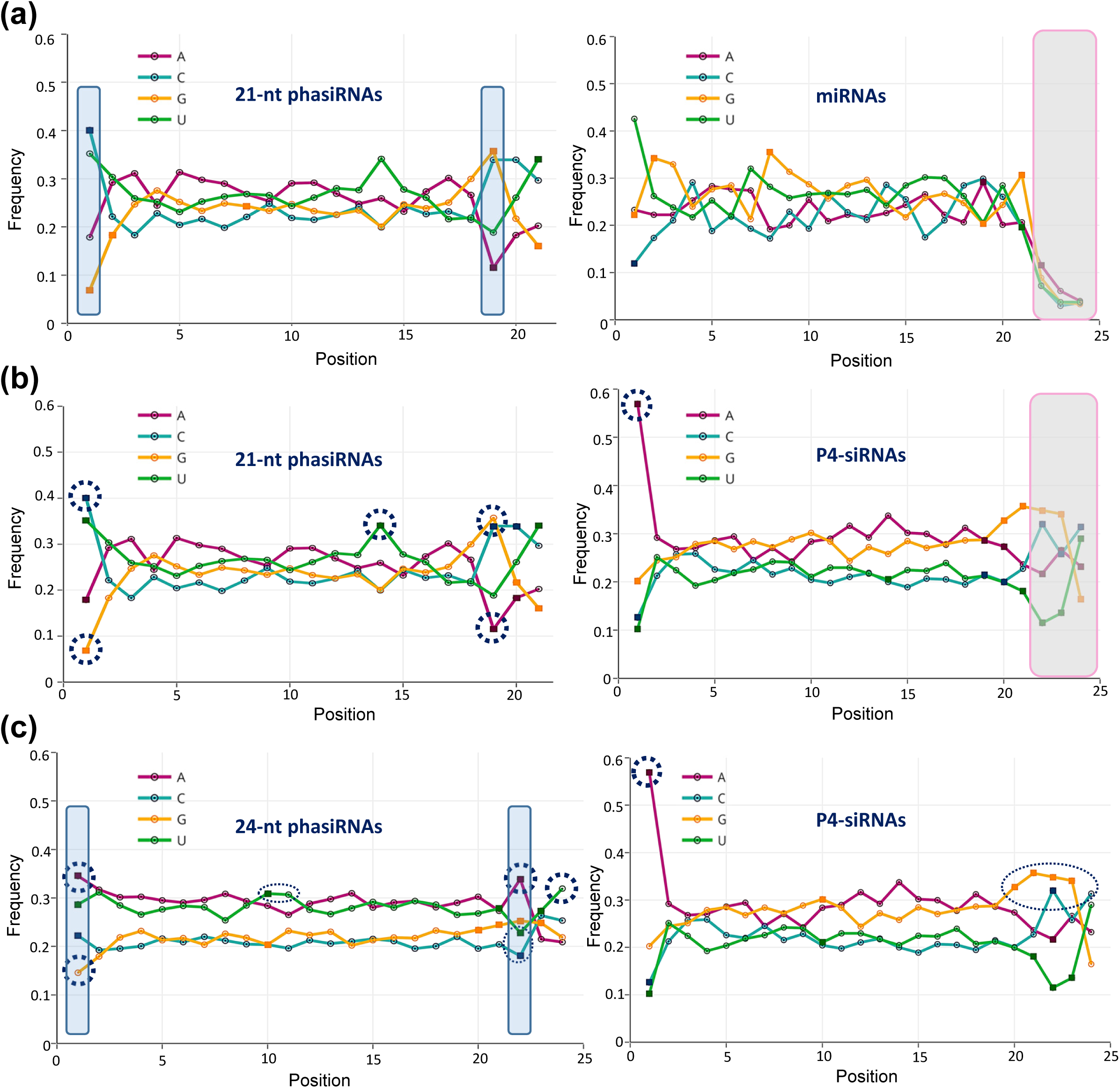
Reproductive phasiRNAs have characteristic position-specific nucleotide biases. Single-nucleotide sequence profiles of position specific base usage comparing 21-nt phasiRNAs (left) and either miRNAs (at right in panel (a)), or 24-nt P4-siRNAs (at right in panel (b)). For all phasiRNA analyses in this figure, the top most abundant 1000 phasiRNAs from the rice and maize data were combined; in panel (a), 553 rice and 203 maize miRBase-annotated miRNAs were used (see Method S2). The frequencies of each of the four bases (A, C, G, and U) at each position are indicated as an open circle. Markers denoted as small square boxes represent positions at which a statistically significant (p = 1e-5) base usage distinguishes phasiRNAs and either miRNAs (panel (a)) or P4-siRNAs (panel (b)), determined by comparison of the data in the two plots. Dotted circles highlight positions in the sequences selected for further discussion in the main text. The gray boxes at right covering the 22nd, 23rd, and 24th positions to retain fair comparison with 21-nt phasiRNAs and the longer sequences, including that those additional position could be disregarded. (c) Single-nucleotide sequence profiles of position specific base usage comparing 24-nt phasiRNAs (at left) and 24-nt P4-siRNAs (at right). In panels (a,c), the blue boxes highlight positions that were analyzed in greater detail in Fig. 4 (positions #1 & 19 for 21-nt phasiRNAs, and positions #1 & 22 for 24-nt phasiRNAs).

At a significance level of 10^−5^, comparing the 21-nt phasiRNAs and miRNAs, we found that the usage of bases at eight positions differed significantly (positions 1, 2, 8, 19, and 21; Fig. 2a). Next, we repeated the calculation, comparing 21-nt reproductive phasiRNAs and 24-nt P4-siRNAs (Fig. 2b), demonstrating significant differences at positions 1, 14, 19, 20, and 21. Combining these results, we made several observations: (i) in these abundant 21-phasiRNAs, there was a 5’ nucleotide preference for C, consistent with a recent report (Komiya et al., 2014), but a strong depletion of G. (ii) We noticed a peak of U at the 14th position in the phasiRNAs (relative to P4-siRNAs), unusual as there were no other biased positions between 3 and 19; the only other internal position showing bias was a G at position 8 in the miRNAs (Fig. 2a). (iii) In the 3’ end of the 21-nt phasiRNAs, we observed a peak of G at the 19^th^ position (with a depletion of A), and U at the 21^st^ position (G strongly disfavored). This representation of G at the 19^th^ position was investigated in more detail below.

We conducted a similar analysis comparing the position-specific base usage between the 24-nt reproductive phasiRNAs and P4-siRNAs. We found that positions 1, 10, 20, 21, 22, and 23 were statistically different (Fig. 2c); in other words, the 24-nt phasiRNAs and P4-siRNAs differed substantially in their base usage over the full length of the molecules. All of the over-represented nucleotides in 24-nt phasiRNAs were either A or U (Fig. 2c); the 5’- and 3’-ends showed differences in the two classes of molecules, and internal positions 10 and 11 were overrepresented for U in the 24-phasiRNAs. These correspond to the same two internal positions critical for directing cleavage by AGO proteins in the case of miRNAs (Carrington & Ambros, 2003), so we noted this for subsequent phasiRNA target analysis (see below). The 3’-end difference was most strikingin the P4-siRNAs, there was a high frequency of G from the 20^th^ to 24^th^ positions and a coincident depletion of U (Fig. 2c), whereas 24-nt phasiRNAs had an overrepresented A at the 22^nd^ position and U at the 3’ end. Therefore, we identified several notable sequence-based features of both classes of reproductive phasiRNAs, observed at both the 5’- and 3’-ends and a small number of internal positions; the 24-nt phasiRNAs also displayed an overall nucleotide composition distinct from that of P4-siRNAs. These differences likely have implications for AGO loading and phasiRNA-target interactions, while also potentially explaining the non-stoichiometric abundances of individual phasiRNAs at each *PHAS* locus.

As observed for animal miRNAs (Chatterjee *et al.*, 2011; Tamim *et al.*, 2018), it’s possible that the non-stoichiometric abundances at a *PHAS* locus results from AGO loading and subsequent stabilization of functional siRNAs. We next computed the sequence profile of ‘present’ or ‘absent’ reproductive phasiRNAs in rice and maize; in other words, at a given *PHAS* locus, some phasiRNAs are never observed in the sequenced sRNAs, but we could extract these computationally and assess their sequence composition biases relative to those we detected experimentally. In a comparison to those phasiRNAs detected in the sequencing data, we observed a substantial, overall sequence composition difference for 21-nt phasiRNAs (Fig. 2 a,c versus Fig. S2a, left). The differences for present versus absent 24-nt phasiRNAs were less pronounced and mainly towards the 3’-end (Fig. 2c, left, versus Fig. S2a, right). To ensure that the profiles for detected phasiRNAs were not unduly biased by the selected use of only the top 1000 sequences, we also plotted sequence profile of all sequenced 21-nt phasiRNAs (Fig. S2b, left) and the 24-nt phasiRNAs (Fig. S2b, right) from the positive set (see Method S2). We observed no noticeable changes in the sequence profile relative to the abundance-selected subset (i.e. Fig. 2 a,c), except a slightly higher representation of 5’U compared to 5’C in the 21-nt phasiRNAs. The comparison of present versus absent reproductive phasiRNAs demonstrated significant differences in nucleotide composition, consistent with relative stabilization of those detected reproductive phasiRNAs after biogenesis; this may reflect AGO loading, target interactions, or other sequence-specific functions of these phasiRNAs.

### The duplex nature of phasiRNA biogenesis impacts nucleotide composition

The observed nucleotide biases at the 19^th^ position in the 21-nt phasiRNAs and at the 22^th^ position in the 24-nt phasiRNAs were the next subject of our investigation. Dicer cleavage of dsRNA typically yields a 2-nt 3’ overhang (Macrae *et al.*, 2006), and thus derived from a long, dsRNA precursor, each sRNA duplex overlaps by two complementary nucleotides at each end, with the neighboring phasiRNAs. In a schematic integrating position-specific biases (Fig. 3a,b), the influence of the most-frequent nucleotides in the “top” strand (the strand generated by RNA polymerase II, which is also targeted by the miRNA trigger) on the composition of the “bottom” strand (the strand generated by RNA DEPENDENT RNA POLYMERASE 6, RDR6) is highlighted for the first and last three nucleotide positions; for example, the 19^th^ position G corresponds to a 5’ C (1^st^ position) for the duplex phasiRNA. Thus, there is a potential co-bias between the 1^st^ and 19^th^ positions, such that if both strands of a 21-nt phasiRNA duplex require a specific 5’ nucleotide to ensure proper AGO loading (like a 5’ C), the 19^th^ position will co-vary with the 1^st^ position. Alternatively, if only one strand of the duplex is loaded (due to a requisite 5’ nucleotide, the primary biogenesis strand, or other reasons) and the duplex partner is dispensable, then the 19^th^ position of the loaded strand is under no selective constraints. For example, in 21-nt phasiRNAs, the 5’ position was predominantly C (40.1%) at the 1^st^ position, and the most prevalent 19^th^ nucleotide was G (35.7%) (Fig. 4a, upper chart). This is consistent with a co-bias for the paired positions in the duplex, yielding duplexes with 5’ C at each end (Fig. 3a). We can infer that 21-nt phasiRNAs may have no strand specificity and either strand is likely to be loaded into the AGO protein as long as there is a 5’ C. Similarly, among the 21-nt phasiRNAs, the 19A and to a lesser extent 19U classes were underrepresented (Fig. 4a, lower), corresponding to bottom-strand 1U and 1A phasiRNAs in a duplex; since 1U phasiRNAs were common among the sequenced phasiRNAs (Fig. 4a, upper), we could infer a bias against 1U phasiRNAs in the complement to phasiRNAs abundant in our libraries.

**Fig. 3.**
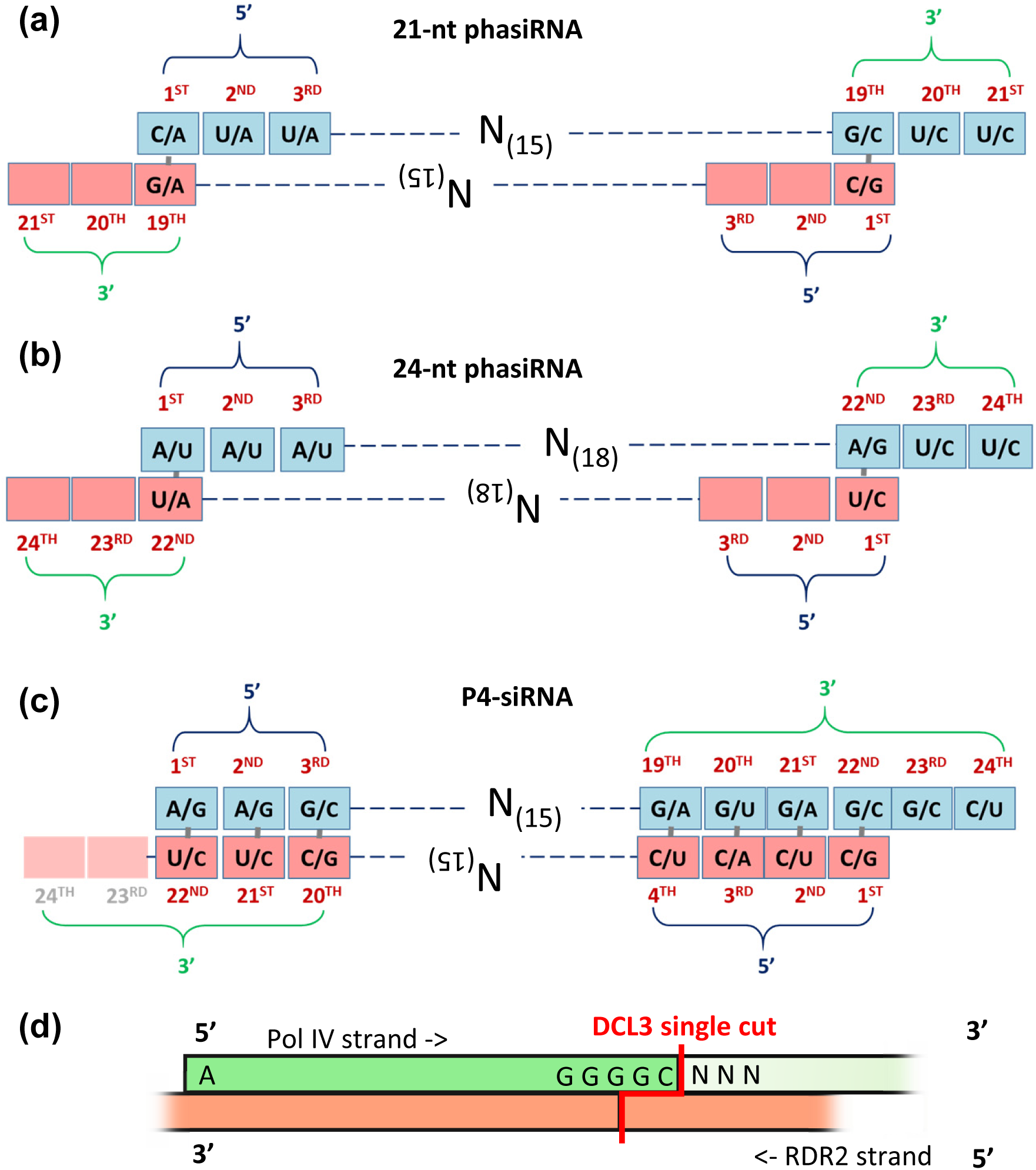
Nucleotide biases indicate one of the two siRNA precursor duplex small RNAs is preferentially retained. Schematic duplex structures of different types of plant small RNAs; the 5’- and 3’-ends are annotated and highlighted to emphasize the influence that a nucleotide bias on one strand has on the other due to pairing. The first three and the last three nucleotide positions are indicated from the 5’- and 3’-end positions, respectively, as the analyses focused on sequence composition biases at these positions; red numbering indicates the base position within the small RNA. Within each position, the top two most frequent nucleotides are indicated, with the first representing the most common occurring nucleotide; the sequences analyzed are the same as Fig. 2. (a) Position-specific nucleotide biases for abundant 21-nt reproductive phasiRNAs in rice and maize. (b) Position-specific nucleotide biases for 24-nt reproductive phasiRNAs from rice and maize. (c) Position-specific nucleotide biases for P4-siRNAs from rice and maize; for P4-siRNAs, the RDR2-derived bottom strand may terminate at the 22^nd^ position, corresponding to the 5’ end of the ‘top’, Pol IV-derived strand, although this is as-yet poorly characterized (indicated by lighter shading of the 23^rd^ and 24^th^ positions). (d) For comparison to panel (c), prior work by Zhai *et al*. (2015) and Blevins *et al*. (2015) described the P4R2 (Pol IV and RDR2-derived) precursors of 24-nt P4-siRNAs as ∼26 to 42 nt RNAs; mapped onto the green Pol IV RNA are the biases observed here for P4-siRNAs. The 5’ and 3’ ends of the RDR2-derived strands are blurred because these ends have not yet been characterized.

**Fig. 4.**
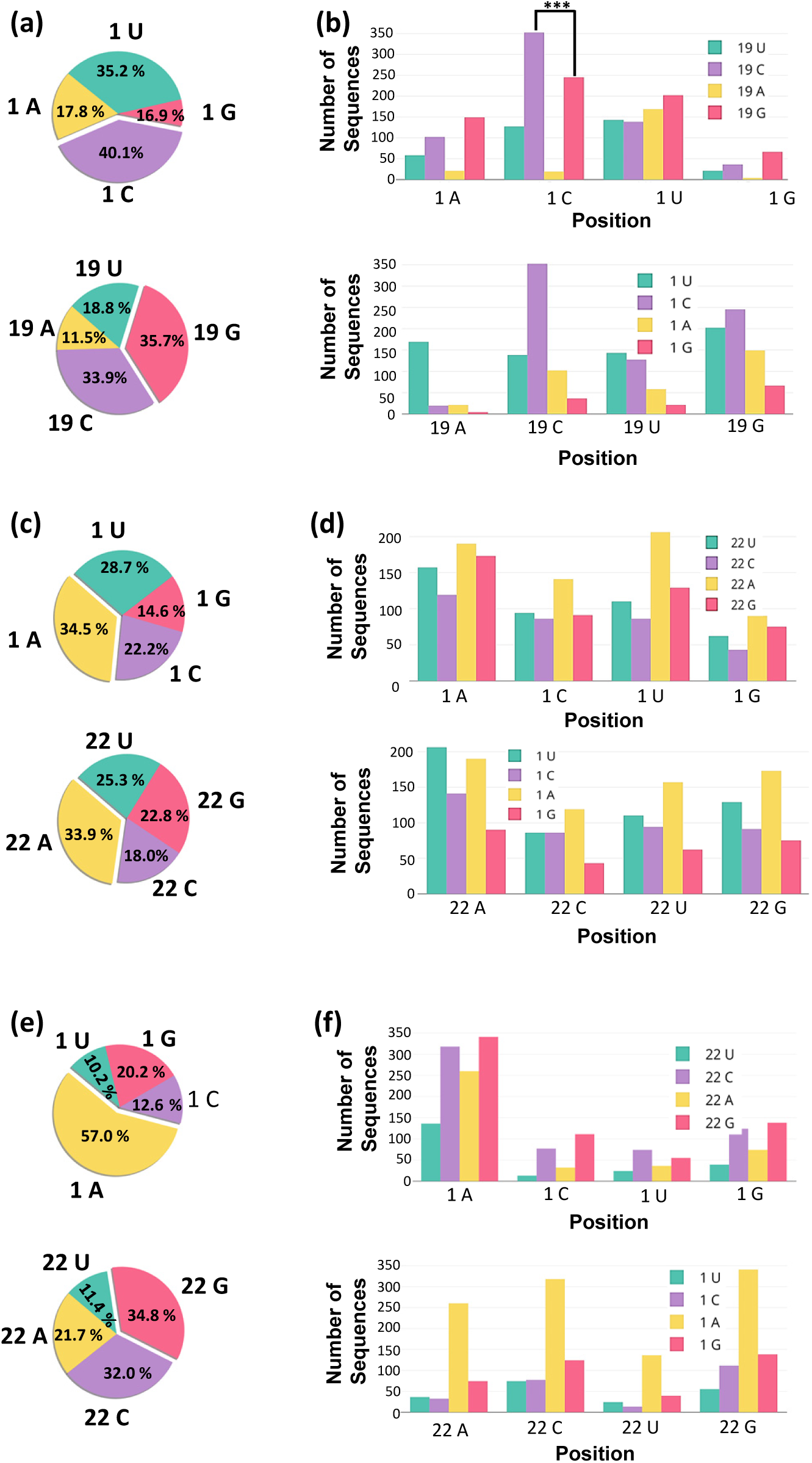
5’ends in phasiRNA duplexes influence the composition of 3’-proximal nucleotides. (a) The pie charts show the composition as a percent of all four nucleotides at the 1st (above) and at the 19th (below) positions in 21-nt reproductive phasiRNAs, combined from maize and rice. The predominant nucleotide is highlighted by separation from the other three. These data are the same as Fig. 2a (blue boxes in that figure), redrawn here for clarity. (b) Above, nucleotide composition at the 19th position of the 21-nt phasiRNAs shown in panel (a) when the 1st position is selected or fixed, as indicated on the X-axis. Below, the same analysis for the 1st position composition when the 19th position is selected or fixed. Significant differences are indicated (Student’s t-test): ***, *P* ≤0.001. (c) Pie charts shows the composition as a percent of all four nucleotides at the 1st and at the 22nd positions in 24-nt phasiRNAs, combined from maize and rice. These data are the same as Fig. 2c, left panel (blue boxes in that figure), redrawn here for clarity. (d) Above, nucleotide composition at the 22nd position of the 24-nt phasiRNAs shown in panel (c) when the 1st position is selected or fixed, as indicated on the X-axis. Below, the same analysis for the 1st position composition when the 22nd position is selected or fixed. (e) Pie charts as above, for P4-siRNAs, combined from maize and rice. These data are the same as Fig. 2c, right panel, redrawn here for clarity. (f) Above, nucleotide composition at the 22nd position of the 24-nt P4-siRNAs shown in panel (e) when the 1st position is selected or fixed, as indicated on the X-axis. Below, the same analysis for the 1st position composition when the 22nd position is selected or fixed.

To assess positional covariance, we analyzed 21- and 24-nt phasiRNAs versus P4-siRNAs, comparing the 5’ nucleotide to the position complementary to the bottom-strand 5’ position (19 in 21-nt siRNAs, and 22 in 24-nt siRNAs). We used these results to make inferences (see the discussion section) about strand specificity in the biogenesis of plant reproductive phasiRNAs. First, we compared the nucleotide composition at the 19^th^ position of 21-nt phasiRNAs for a given 1^st^ nucleotide and we performed the same analysis for the 1^st^ position composition with the 19^th^ position fixed (Fig. 4b). The 1U phasiRNAs (i.e. 5’ U) had an almost uniform distribution of nucleotides at the 19^th^ position, which was striking relative to the 1C, 1A, and 1G phasiRNAs, which were depleted for 19A phasiRNAs (and 19U, to a lesser extent). Another noticeable bias was for 1C phasiRNAs, which were predominantly 19C or 19G, yielding a phasiRNA duplex of either 1C/1G or 1C/1C (top strand/bottom strand). 19G was prevalent for 1A, 1U, and 1G phasiRNAs, which in each case would yield a 1C bottom-strand phasiRNA. Next, we analyzed the 5’ nucleotide composition for 21-nt phasiRNAs after fixing the 19^th^ position (Fig. 4b, lower panel). Among 19G phasiRNAs (the predominant group based on Fig. 4a), 1C was most common, corresponding to a 1C/1C duplex. For 19C phasiRNAs (1G on the complement), a strong bias of 1C was observed; since 1C 21-nt phasiRNAs are most commonly loaded to MEL1 (Komiya *et al.*, 2014), this was perhaps an indication of strand specificity (i.e. 1C/1G duplexes, so only the 1C strand loaded). Therefore, among 21-nt phasiRNAs, there is a co-bias of the 1^st^ and 19^th^ positions, perhaps reflective of strand specificity in AGO loading.

Next, we performed similar analyses for 24-nt phasiRNAs, focused on the 1^st^ and 22^th^ positions (Fig. 3b). The 1^st^ position was less biased than 21-nt phasiRNAs, although 1G was also underrepresented (Fig. 4c, upper); at the 22^nd^ position, there was less bias than for the 19^th^position of the 21-mers (Fig. 4c, lower), with an increase of A representation, particularly relative to other nucleotide positions (Fig. 2c, left). 22A corresponds to 1U in the complement, and since 1U 24-nt phasiRNAs were common in our dataset (Fig. 4c, upper), both phasiRNAs in such a duplex are favored in our data, consistent with a lack of strand specificity. Lower levels of 1^st^/22^th^ position covariation were observed in 24-nt than 21-nt phasiRNAs (Fig. 4d), and there was an overall A-U enrichment (Fig. 2c), demonstrating more relaxed sequence constraints.

For comparison to the 24-nt phasiRNAs, we measured the position-specific nucleotide biases for P4-siRNAs. Their precursors have been described (Fig. 3d; summarized from Blevins *et al.*, 2015; Zhai *et al.*, 2015a), although the nature of RDR2-derived bottom strands is as-yet incompletely understood (i.e. how they initiate and terminate relative to the ends of the P4 precursor). Unlike phasiRNAs, however, there is no expectation of P4-siRNA “duplexes” whereby either strand could be loaded, and data from Zhai *et al.* (2015a) indicate that the P4 strand is preferably loaded over the RDR2 strand (Fig. 3c). Apart from the strong overall 1A bias mentioned above, no notable co-variation biases were observed (Fig. 4e,f); i.e. the proportional representation in the 22^nd^ position was essentially invariant, regardless of the 1^st^ position nucleotide, G>C>A>U, consistent with a strong overall bias to the GGGGC motif in the 3’ end (Fig. 2c).

Combining the compositional analyses described above, we applied these same approaches to an unusual group of siRNAs, a set of 22-nt, putative heterochromatic siRNAs that are RDR2-independent, thus far found only in maize (Nobuta *et al.*, 2008). We were interested to analyze these “22-nt hc-siRNAs” because they are poorly characterized and their relationship to P4-siRNAs is not known (see Method S4 for extracting 22-nt siRNAs). The most significant difference between 22-nt hc-siRNAs and 24-nt P4-siRNAs was at 5’ end positions 1, 3 and 4 (Fig. S3 a,b), but the level of A in 22-nt hc-siRNAs was significantly lower from position 12 to the 3’ end, compared to the 24-nt P4-siRNAs. There were apparent 3’ differences as well, but this was from the comparison performed by counting nucleotides from the 5’ end. We reassessed differences by aligning the 3’ ends and measuring positions starting from the 3’ end (i.e. comparing up to five positions at the 3’ end minus N nucleotides), in case AGO binding occurs in some cases from the 3’ end. Measured this way, we observed only one 3’ difference, at the 3’ end – 1 position, at which the G-U composition varied significantly (Fig. S3c). We next looked at covariation between the 20^th^ and 1^st^ nucleotides in the 22-nt hc-siRNAs; as with P4-siRNAs, the 20^th^ nucleotide representation was more or less the same for all 5’ nucleotides, and even for the major class of 5’ U siRNAs, 20^th^ position G or C nucleotides were equally represented (Fig. S3d). This lack of bias would yield many bottom strand 5’ G sRNAs which are disfavored, consistent with strand specificity for the 22-nt hc-siRNAs (Fig. S3e). Thus, these RDR2-independent 22-nt siRNAs may be produced by the activity of other RDRs such as RDR1 or RDR6; although the RNA polymerase generating their primary strand precursor remains to be determined, the 5’ difference of 22-nt hc-siRNAs compared to P4-siRNAs suggests an alternative production pathway and/or function.

The results of analysis of the nucleotide and co-variation biases across different classes of siRNAs at the 5’ and 3’-proximal ends are consistent with evidence of strand specificity for both 21- and 24-nt phasiRNA duplexes. There is stronger support for strand selection of 21-nt reproductive phasiRNAs, perhaps reflective of selection by the AGO protein of one strand over the other.

### Predicted targets of reproductive phasiRNAs as a means to infer function

As little is known about the targets and the functions of the reproductive phasiRNAs, we attempted to predict targets for the 500 most abundant pre-meiotic (21-nt) and meiotic (24-nt) phasiRNAs in rice. Using standard criteria (i.e. modeled on known miRNA-target interactions), prior reports have failed to find targets of reproductive phasiRNAs, while reporting few details of these analyses due to the negative result (Song *et al.*, 2012b; Zhai *et al.*, 2015b). We revisited this topic because new, more powerful, faster and flexible target prediction methods are available; prior work used a “seed-based” sRNA-target interaction pipeline, which is derived from models of animal miRNAs and does not accurately capture the target similarity of most plant miRNAs (Kakrana *et al.*, 2014). We used sPARTA (Kakrana *et al.*, 2014) based on a “seed-free” approach and allows greater flexibility in pairing parameters. To gain insights about phasiRNA targeting, we conducted a comparative analysis, measuring class-by-class how predicted targets of these abundant phasiRNAs compared to those of other known sRNAs, such as miRNAs and P4-siRNAs.

#### 21-nt phasiRNAs

First, we compared in rice the distribution of predicted target scores (TS) of 21-nt phasiRNAs with a selected set of known, conserved miRNAs (Fig. 5a). We selected plant miRNAs with numbers lower than miR1000 (i.e. osa-miR162) (n=288), as these are generally abundant, conserved, and better characterized than any more recently-described miRNAs. For each class, miRNAs versus 21-nt phasiRNAs, targets were predicted using sPARTA (Kakrana *et al.*, 2014). We retained two sets of results, either all targets or only the “best” targets (those with a lowest target penalty score, meaning a high degree of complementarity). Each sRNA would also have at least one perfect match in the genome, a target score of 0, potentially the result of targeting in *cis*. For 21-nt phasiRNAs, the TS distribution showed a peak in the number of best targets at 3.5 (Fig. 5a, left), compared to ∼1 for miRNAs (Fig. 5a, right). The relative paucity of TS matches in the range of 0.5 to 1.5 for 21-nt phasiRNAs was striking, particularly since many miRNAs have predicted targets in this range. We inferred based on this pattern of sequence complementarity that 21-nt phasiRNAs, unlike miRNAs, either may function largely in *cis* via perfect matches or have been selected to avoid closely-matched targets.

**Fig. 5.**
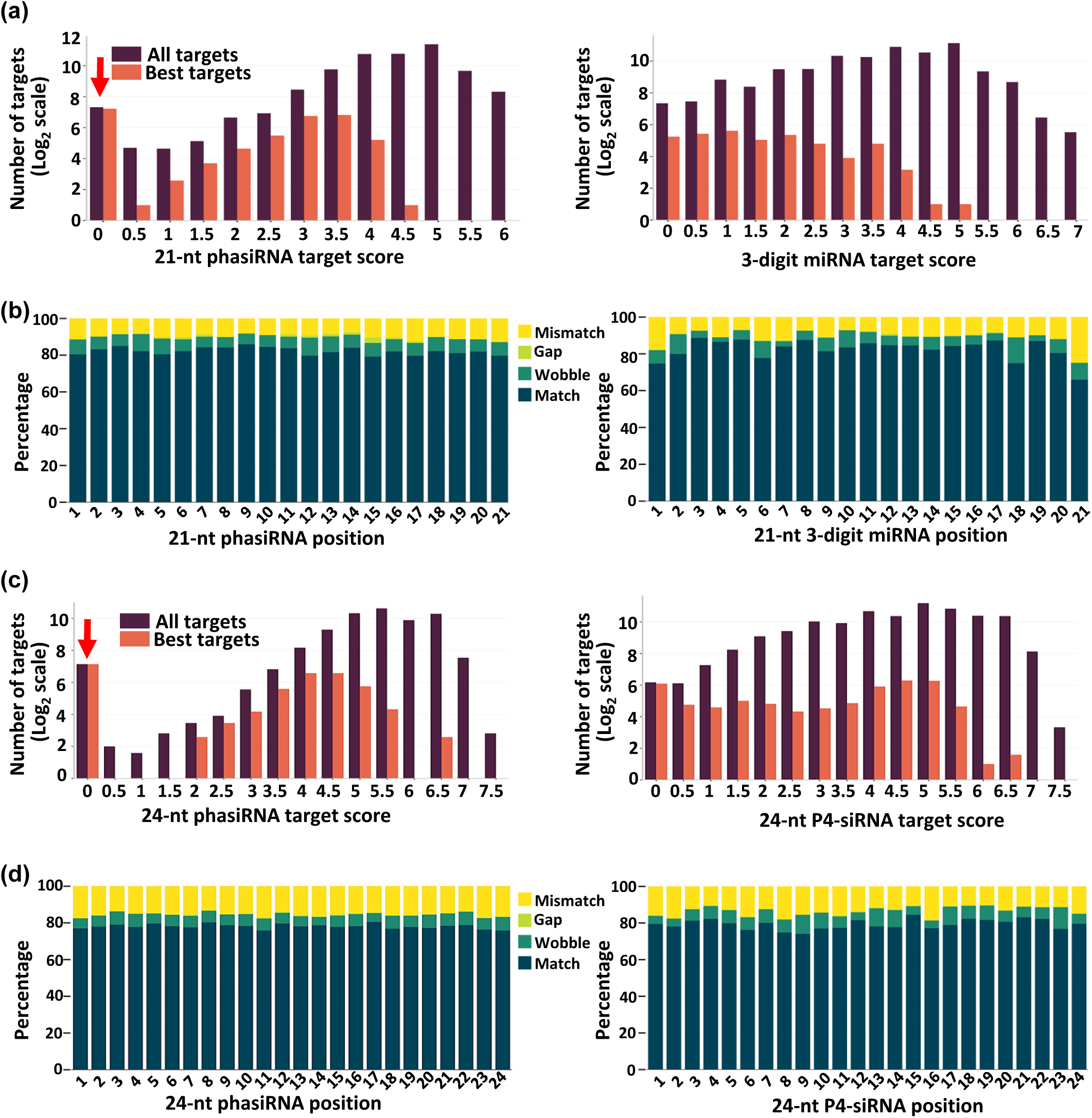
phasiRNA target prediction illustrates low binding affinity compared to other sRNAs to their targets due to sequence diversity. Target prediction for top 500 most abundant 21- and 24-nt phasiRNAs in rice, rice 3-digit miRNAs (n=288), and top 500 most abundant P4-siRNAs in rice was performed using sPARTA. (a) The bar plots show target score distribution (as indicated on X-axis) for 21-nt phasiRNAs (at left) and 3-digit miRNAs (at right). Dark purple bars depict target score distribution of all targets of 21-nt phasiRNAs and 3-digit miRNAs. Orange bars depict target score distribution of only best targets (targets with a lowest target penalty score, meaning high degree of complementarity between phasiRNAs or miRNAs and their targets) of 21-nt phasiRNAs and 3-digit miRNAs. As indicated, Y-axis (number of targets) is transformed into log2 scale and red arrow indicates potential self-targeting or *cis* interactions (with target score of 0, meaning perfect match). (b) The bar charts record the 21-nt phasiRNA-target interaction (at left) and 3-digit miRNA-target interaction (at right) for all targets with target score between 0.5 and 3.5, capturing binding pattern as a percent (Y-axis) of match, gap, wobble, and mismatch. (c) Bar plots showing target score distribution as above panel (a), for 24-nt phasiRNAs (at left) and 24-nt P4-siRNAs (at right). (d) As above panel (b), the bar charts indicating the binding pattern as a percent (Y-axis) of match, gap, wobble, and mismatch for 24-nt phasiRNA-target interaction (at left) and 24-nt P4-siRNAs-target interaction (at right) for all targets with target score between 0.5 and 5.

To dissect these predicted sRNA-target interactions in rice, we recorded position-specific matches for both 21-nt phasiRNA-target interactions and 21-nt miRNA-target interactions (Fig. 5b). This represented the putative binding pattern as a percentage of each position of predicted matches, gaps, wobbles, and mismatches. We selected only predicted targets (for both phasiRNAs and miRNAs) with a TS between 0.5 and 3.5, omitting self-targeting interactions. Overall, consistent with higher scores, 21-nt phasiRNAs showed lower match rates across all positions than miRNAs (Fig. 5b); a few substantial position-specific differences were observed, including higher match rates for phasiRNAs at the 1^st^ and 21^st^ positions, and a higher (yet unexplainable) rate of gaps at the 15^th^ position (Fig. 5b). We concluded that unless 21-nt reproductive phasiRNAs target primarily in *cis,* they must have lower levels of complementarity to their targets than miRNAs.

#### 24-nt phasiRNAs

Next, we extended our analysis to attempt to find the targets of the reproductive 24-nt phasiRNAs, again focusing on rice. We performed similar analyses as above and compared the TS distribution of 24-nt phasiRNAs (Fig. 5c, left) with the top 500 most abundant 24-nt P4-siRNAs (Fig. 5c, right). For the 24-mers, we omitted the higher penalty for a mismatch at the 10^th^ and 11^st^ positions in the target alignment; that penalty is relevant for 21/22-nt sRNAs that direct cleavage at those positions, whereas pairing requirements for individual 24-nt siRNAs have not been described or tested. For the 24-nt phasiRNAs, we observed a peak in the number of best targets at 4.5 (Fig. 5c, left); while score of 4.5 to 5 was also the peak for P4-siRNAs (excluding perfect, or ‘cis’ matches at 0), P4-siRNAs had a much more even distribution of scores. There was a striking gap in the distribution of target scores from 0 to ∼2 for the 24-nt phasiRNAs, indicating that these lack highly homologous *trans* targets (Fig. 5c, left). In other words, the 24-nt phasiRNAs are largely quite distinct from most other genome sequences, relative to P4-siRNAs, which find many highly homologous potential target sites.

Again, as for the 21-nt phasiRNAs, we predicted and recorded position-specific matches for both 24-nt phasiRNA-target interactions and P4-siRNA-target interactions (Fig. 5d). This represented the putative binding pattern as a percentage of each position of predicted matches, gaps, wobbles, and mismatches. In this case, given the different score distribution relative to 21-mers, we selected only predicted targets (for both phasiRNAs and P4-siRNAs) with a TS between 0.5 and 5, omitting self-targeting interactions. Overall, consistent with higher TS scores, 24-nt phasiRNAs showed much lower match rates across all positions than P4-siRNAs (Fig. 5d left versus right), i.e. an average of 15 to 20% mismatches compared to fewer than 15% mismatches for 24-nt P4-siRNAs.

##### Classes of predicted reproductive phasiRNA targets

As a final step in analyzing the possible targets of reproductive phasiRNAs in rice, we classified the predicted target loci. This analysis used all predicted targets described in the sections above, including both *cis* and *trans* targets. In rice, the top 500 21-nt phasiRNAs were predicted to target 7766 loci (Table S2). These putative targets included 1400 (18.02 percent) loci classified by RepeatMasker as related to the transposable elements (TEs). The top 500 24-nt phasiRNAs were predicted to target 5631 loci, of which 836 (14.84 percent) are related to TEs (Table S3). To assess whether these predicted matches to TEs represent an enrichment or depletion compared to random chance, we randomly selected 7800 and 5600 genes from the 35,000+ annotated genes in rice; among these, ∼30 to 31% are TE-like. Therefore, the predicted targets of reproductive phasiRNAs are relatively depleted for TE-like targets. Overall, our more detailed results are consistent with earlier statements that classes of potential targets are not evident for reproductive phasiRNAs, and thus the characterization of their functions will require molecular and biochemical investigation.

## Discussion

Our machine learning-based workflow focused on sequenced-based and structural features of plant sRNAs, with an emphasis on the poorly characterized set of reproductive phasiRNAs. We demonstrate that this approach can successfully classify reproductive phasiRNAs relative to other endogenous plant sRNAs, with high values for ACC, SE, SP, PPV, and AUC. Feature selection demonstrated the importance of the 5’- and 3’-ends, k-mer features, GC content, and structural features including the MFE. We observed characteristics that may reflect specificity in AGO loading of reproductive phasiRNAs, the key to the function of all sRNAs. Examination of spatiotemporal expression data for AGOs in rice and maize shows a high correlation between peaks of abundance of reproductive phasiRNAs and *AGO* genes, suggesting that there might be a functional connection. From rice and maize data, this includes OsAGO1d, ZmAGO18b, OsAGO18, OsAGO2b (Zhai *et al.*, 2015b; Fei *et al.*, 2016), and OsAGO5c (MEL1) which loads 21-nt phasiRNAs in rice. In Arabidopsis, AGO3, close to OsAGO2b (Zhang *et al.*, 2015), recruits 24-nt sRNAs with 5’A and effects epigenetic silencing, consistent with the hypothesis that 5’A 24-nt phasiRNAs might be loaded into AGO2b in grasses. Moreover, ZmAGO18b, a grass specific AGO, binds both 21-nt phasiRNAs with 5⍰U and 24-nt phasiRNAs with 5⍰A to function in inflorescence meristem and tassel development (Sun *et al.*, 2017). Our classification data lay the groundwork for better definition of AGO-phasiRNA interactions.

One unique aspect of working with the reproductive phasiRNAs is that their production from long, double-stranded RNA precursors from hundreds or thousands of loci yields a rich dataset for which comparable analyses of tasiRNAs or miRNAs are not possible due to their more limited representation. This allowed the large-scale assessment of biases in representation in the libraries, from which we observed significant biases in the representation of specific nucleotides at the 1^st^ and 19^th^ positions among the 21-mers. One possible interpretation of these biases is a model of competition for loading between the two strands of a duplex, whereby one strand is preferentially loaded over the other, typically understood to be driven by the 5’ nucleotide (Schwarz *et al.*, 2003), which is a preferred C in the case of 21-nt reproductive phasiRNAs (Komiya *et al.*, 2014). Yet, 1U phasiRNAs are quite abundant, begging the question of whether these are competing with 1C phasiRNAs for loading into MEL1; among sequenced MEL1-associated phasiRNAs, 1U phasiRNAs were less than 10% of the total (Komiya *et al.*, 2014). Perhaps the higher proportion in the sequenced phasiRNAs reflects (1) stability in the absence of loading, or (2) perhaps 1U phasiRNAs are loaded into a different AGO than the 1C phasiRNAs – maybe AGO1, known to have an affinity for 1U 21-nt sRNAs (Zhao *et al.*, 2016). Assuming the latter, for the sake of argument, the difference in the 19^th^ position for a given 1^st^ position nucleotide for the 21-nt reproductive phasiRNAs could be explained by AGO affinity: 1U phasiRNAs may be loaded as well or better than 1C phasiRNAs, but into this different AGO. An additional influence on these terminal or near-terminal positions may be strand selection during AGO loading of the duplex, which is influenced by factors including the thermodynamic stability of the two ends of each phasiRNA duplex (Schwarz *et al.*, 2003).

Based on the observation of abundant 1U and 1C 21-nt phasiRNAs, we hypothesized an AGO competition model (Fig. S4). We inferred/hypothesized this because of the data in Fig. 4B (upper panel) that the sequenced 1V (V = A or C or G, using the IUPAC code) phasiRNAs are depleted for 19A phasiRNAs, which would be 1U on the bottom strand; perhaps this is because in a duplex with a 1U phasiRNA, the 1U phasiRNA is loaded. But sequenced 1U phasiRNAs showed no bias in the 19^th^ position, because they are preferred over the opposite strand, and thus are the “winners” in the competition (Fig. S4a). In contrast, the 1R/19G (R = A or G) phasiRNAs are paired with 1C phasiRNAs, which is AGO loaded (Fig. S4b). The 1V/19C phasiRNAs are abundant because these are paired with 1G phasiRNAs, which are not AGO loaded and thus are always “losers” in the competition with their duplex pairs. The 1C phasiRNAs are an interesting case because based on frequency, 1C/19C > 1C/19G > 1C/19W (W = A or U) (Fig. 4b, upper panel). The 1C/19G phasiRNAs are paired with 1C phasiRNAs, which compete well, and thus either strand may be loaded and stabilized (Fig. S4c). The 1C/19U phasiRNAs are less frequent because they are paired with 1A phasiRNAs that are not particularly stabilized or loaded. In other cases (Fig. S4d), the frequency of 1D/19G phasiRNA is higher than 1D/19C (D = A or G or U) (Fig. 4b, upper panel); one interpretation of the high frequency of 1D/19G is that since the 1D/19G phasiRNAs are paired with a 1C/19H phasiRNA (H = A or C or T), thus 1C/19H phasiRNA is preferentially loaded and stabilized. Thus, phasiRNAs from a 1D/19G duplex are more abundant than those from a 1D/19C duplex because in the latter, the 1G/19H phasiRNA on the bottom strand is likely not loaded or stabilized. With as rich a dataset as reproductive phasiRNAs provide, we can start to resolve the sequence-based characteristics that influence representation in sequencing data, and infer the mechanistic basis for these differences. For example, we identified novel position-specific biases, like the 14^th^ position in the 21-nt phasiRNAs (Fig. 2a, left, and Fig. S2b, left). These internal positions may be important for AGO loading, or phasing function/targeting, and thus future functional or structural studies should investigate these in greater detail.

## Acknowledgments

We are grateful to members of the Meyers lab for useful discussions. We also acknowledge the contribution of maize *mop1 (rdr2)* mutant tissue from the lab of Vicki Chandler, and Stacey Simon for the construction of those libraries.

## Author Contributions

Experiments were designed by P.P., H.S., S.M., and B.C.M. P.P. implemented methods and conducted the analyses. A.K. contributed methods and algorithmic refinements. H.S. contributed conceptual ideas. S.M. performed data generation. P.P. and B.C.M. wrote the paper with input from all authors; all authors read and approved the manuscript.

## Supplemental Information

The following materials are available in the online version of this article.

**Fig. S1** Information gain (IG) based feature selection

**Fig. S2** Sequence profiles of absent phasiRNAs and all detected phasiRNAs from *PHAS* loci

**Fig. S3** 22-nt siRNAs from maize are distinct from P4-siRNAs

**Fig. S4** An AGO competition model

**Table S1** sRNA libraries from maize, rice and *Setaria viridis* used in this study

**Table S2** Predicted targets of 21-nt phasiRNAs in rice

**Table S3** Predicted targets of 24-nt phasiRNAs in rice

**Table S4** Top 30 features, from example comparisons, obtained using information gain.

**Method S1** Dataset used for cross validation study

**Method S2** Features included in the machine learning algorithm, and their selection

**Method S3** Computational analysis of sequencing data

**Method S4** Extraction of a set of maize 22-nt hc-siRNAs

