## Supplementary Materials for "Reproductive phasiRNAs in grasses are compositionally distinct from other classes of small RNAs"

The following Supporting Information is available for this article:

**Fig. S1** Information gain (IG) based feature selection

**Fig. S2** Sequence profiles of absent phasiRNAs and all detected phasiRNAs from *PHAS* loci

**Fig. S3** 22-nt siRNAs from maize are distinct from P4-siRNAs

**Fig. S4** An AGO competition model

**Table S1** sRNA libraries from maize, rice, and *Setaria viridis* used in this study

**Table S2** Predicted targets of 21-nt phasiRNAs in rice

**Table S3** Predicted targets of 24-nt phasiRNAs in rice

**Table S4** Top 30 features, from example comparisons, obtained using information gain.

**Method S1** Dataset used for cross validation study

**Method S2** Features included in the machine learning algorithm, and their selection

**Method S3** Computational analysis of sequencing data

**Method S4** Extraction of a set of maize 22-nt hc-siRNAs

**Fig. S1 Information gain (IG) was used to reduce feature dimensionality together with ROC curve to select top most 250 most informative features.**
Strip plot shows the distribution of all 1498 features computed for the classification of 21- and 24-nt phasiRNAs. The X-axis shows IG computed for each feature. Features are ranked from high IG to low IG; Y-axis shows the category to which each feature belongs (see Method S1 for more details on k-mers). The dotted gray line represents 250th feature (random cutoff) used for selecting features for classification. Therefore, all features to the left side of dotted line were used in the classification. (a) The 250 features selected based on IG for classification of the 21-nt phasiRNA vs. the set of miRNAs + P4-siRNAs + tRNAs + rRNAs. (b) The 250 features selected based on IG for classification of the 24-nt phasiRNA vs. the set of miRNAs + P4-siRNAs + tRNAs + rRNAs. (c) ROC curve for the classifier used in panel (a), demonstrating an area under the curve (AUC) of 0.97; solid lines show the AUC for each fold up to the 5-fold cross-validation ("CV"). The black dotted line depicts random chance (0.5, diagonal); the blue line shows mean of AUC of for 5-fold CV with SD. Red shaded area represents ±1 SD. The X-axis is plotted as 1 minus the specificity. (d) As in panel C, ROC curve for the classifier used in panel B, demonstrating a mean AUC of 0.942 for 5-fold CV. (e) Accuracy (Y-axis) of the top IG-based and ranked features (on the X-axis) used in panel a (dark gray) and those used in panel (b) (blue).

Fig. S1

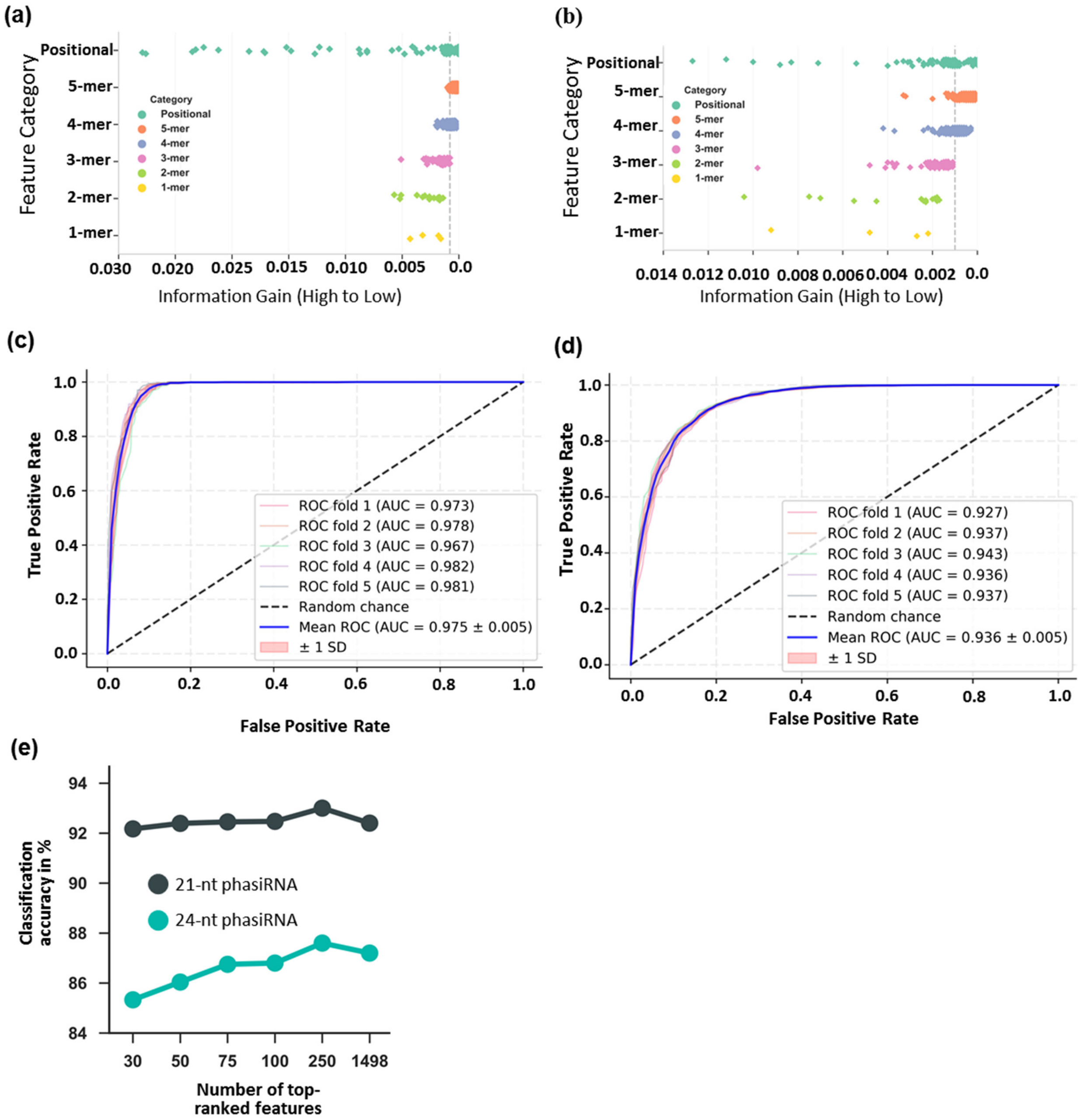
(a)
Feature Category
Positional
5-mer
4-mer
3-mer
2-mer
1-mer
Category
Positional
5-mer
4-mer
3-mer
2-mer
1-mer
0.030
0.020
0.025
0.015
0.010
0.005
0.0
Information Gain (High to Low)
(b)
Feature Category
Positional
5-mer
4-mer
3-mer
2-mer
1-mer
Category
Positional
5-mer
4-mer
3-mer
2-mer
1-mer
0.014
0.012
0.010
0.008
0.006
0.004
0.002
0.0
Information Gain (High to Low)
(c)
True Positive Rate
False Positive Rate
ROC fold 1 (AUC = 0.973)
ROC fold 2 (AUC = 0.978)
ROC fold 3 (AUC = 0.967)
ROC fold 4 (AUC = 0.982)
ROC fold 5 (AUC = 0.981)
Random chance
Mean ROC (AUC = 0.975 ± 0.005)
± 1 SD
(d)
True Positive Rate
False Positive Rate
ROC fold 1 (AUC = 0.927)
ROC fold 2 (AUC = 0.937)
ROC fold 3 (AUC = 0.943)
ROC fold 4 (AUC = 0.936)
ROC fold 5 (AUC = 0.937)
Random chance
Mean ROC (AUC = 0.936 ± 0.005)
± 1 SD
(e)
Classification accuracy in %
21-nt phasiRNA
24-nt phasiRNA
30
50
75
100
250
1498
Number of top-ranked features

**Fig. S2 Position-specific nucleotide biases in either the not-detected positions from reproductive *PHAS* loci phasiRNAs or in all detected phasiRNAs.**
(a) Single-nucleotide sequence profile of position-specific base usage of phasiRNA positions not detected in the sequencing data but within reproductive PHAS loci. At left, 21-nt phasiRNAs (rice = 72955 and maize = 2085, combined); at right, 24-nt phasiRNAs (rice = 1877 and maize = 3916, combined). The frequencies of each of the four bases (A, C, G, and U) at each position are indicated as an open circle. (b) As in panel (a), single-nucleotide sequence profile of position-specific base usage, in this case, for all reproductive phasiRNAs detected in the sRNA libraries used in this study; at left, 21-nt phasiRNAs (rice = 20240 and maize = 9260), and at right, 24-nt phasiRNAs (rice = 2950 and maize = 8800).

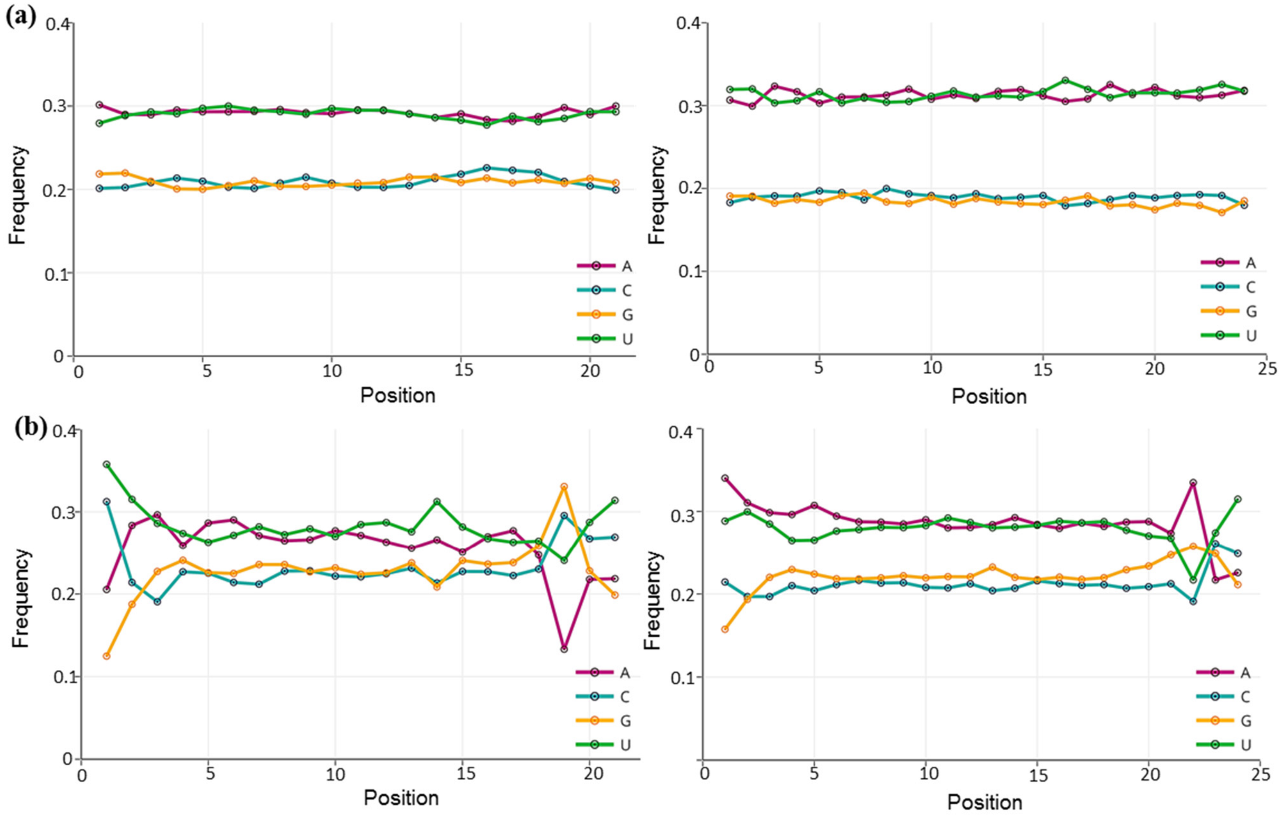

**Fig. S3 22-nt siRNAs from maize are distinct from P4-siRNAs, but have similar 3' end sequence composition.**

(a-b) Single-nucleotide sequence profiles of position specific base usage comparing maize 22-nt siRNAs (N~4000) enriched in libraries from the maize *rdr2* mutant and overlapping with highly repetitive regions (i.e. hits > 50 in the genome) (a) and 24-nt P4-siRNAs, combined from maize and rice (b). The blue boxes highlight positions that were analyzed in greater detail (positions 1 & 20 for 22-nt siRNAs). (c) A comparison of the position-specific base usage of 3'-most five nucleotides of the 22-nt siRNA (left) and P4-siRNAs (at right); for this analysis the small RNAs were aligned from the 3' ends. These five positions were denoted, counting backward from the 3' end; as "3' end", "3' end – 1", "3' end – 2", "3' end – 3", and "3' end – 4". (d) At left, pie charts indicate the composition as a percent of all four nucleotides at the 1st and at the 20th positions in 22-nt siRNAs. These data are the same as panel (a) (blue boxes), redrawn here for clarity. At right, and above, nucleotide composition at the 20th position of the 22-nt siRNAs shown in panel (d) when the 1st position is selected or fixed, as indicated on the X-axis. At right, below, the same analysis for the 1st position composition when the 20th position is selected or fixed. Significant differences are indicated (Student's t-test): ns, not significant. (e) A schematic indicating the top and bottom strand position-specific nucleotide biases for 22-nt siRNAs from maize, as in Fig. 3.

**Fig. S3**

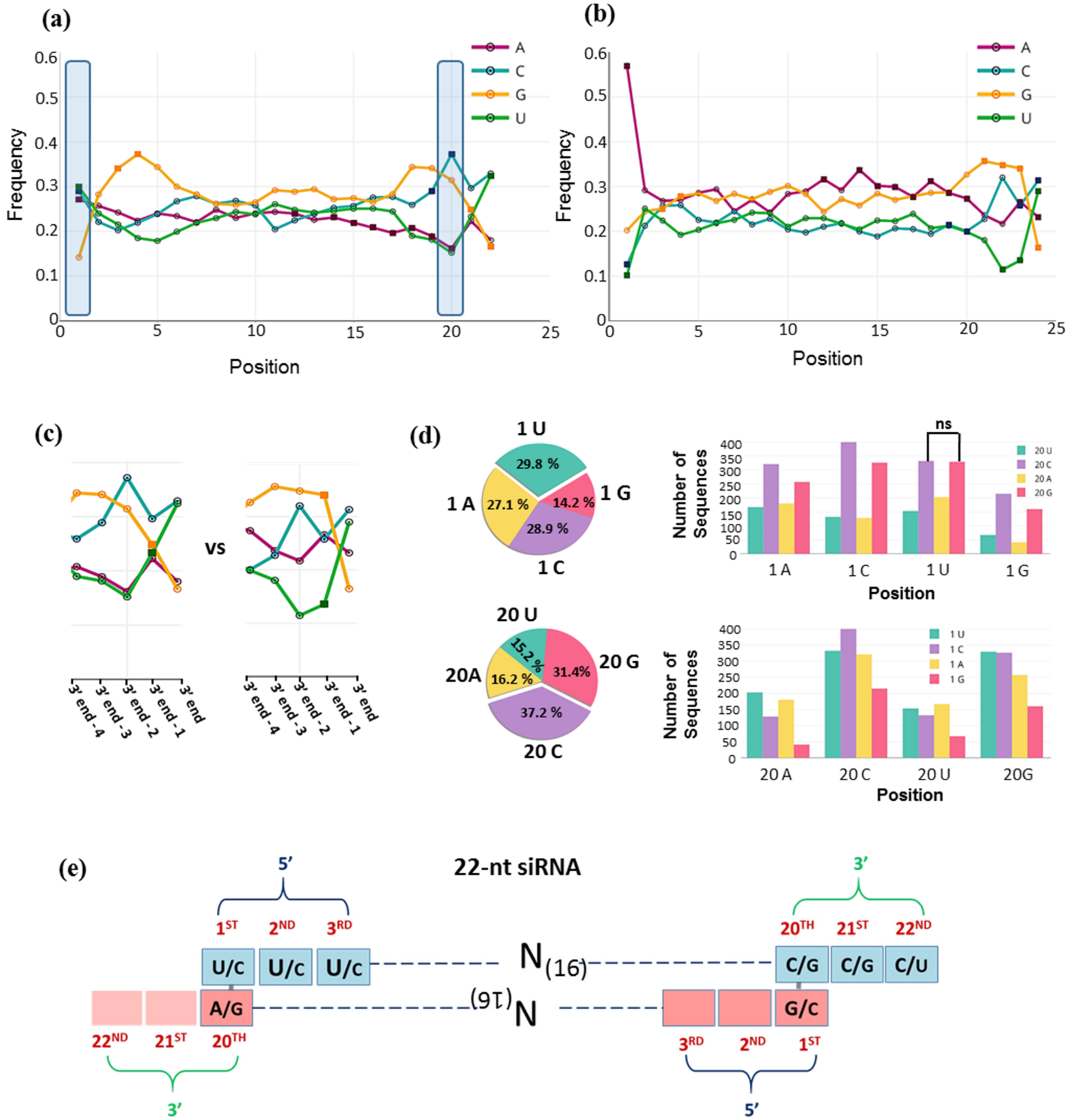
(a)
(b)
Frequency
Position
A
C
G
U
(c)
vs
3' end - 4
3' end - 3
3' end - 2
3' end - 1
3' end
(d)
1 U
29.8 %
1 G
14.2 %
1 A
27.1 %
28.9 %
1 C
20 U
15.2 %
20 G
31.4 %
20A
16.2 %
37.2 %
20 C
ns
Number of Sequences
1 A
1 C
1 U
1 G
20 A
20 C
20 U
20G
(e)
22-nt siRNA
5'
3'
1ST
2ND
3RD
20TH
21ST
22ND
U/C
C/G
C/U
A/G
G/C
N(16)

**Fig. S4 An AGO competition model may explain abundance differences for 1U, 1C or other 21-nt phasiRNAs.**

We propose a model for AGO proteins competing for phasiRNAs, distinguished by 1st and 19th position nucleotides, to explain the substantial differences in abundances observed among sequenced or absent phasiRNAs, and when compared to MEL1-associated 21-nt phasiRNAs. IUPAC nucleotide codes (shown in the table) are used for simplifying the representation of nucleotides at the 1st and the 19th positions, highlighted in 'red', in an orange top strand and a gray bottom or complementary strand. Different combinations of the nucleotides at the 1st and the 19th positions on a top and a bottom strand, comprising a single duplex are denoted in "pink" or "purple". Loading of a particular strand from a duplex into a specific AGO is indicated by the arrow of same color as the nucleotides on that strand. (a) In a 1V/19A - 1U/19B duplex, 1U/19B (bottom) phasiRNA is more likely to be loaded into AGO1. In a 1U/19N – 1N/19A duplex, 1U/19N (top) is more likely to be loaded into AGO1. (b) As in panel (a), in a 1R/19G – 1C/19Y duplex, 1C/19Y is preferentially loaded into MEL1. In a 1V/19C - 1G/19B duplex, the 1V/19G phasiRNA is either loaded into MEL1 or another AGO (AGO2?). The 1G/19C phasiRNA is likely not loaded into AGO. (c) In a 1C/19G - 1C/19G duplex, phasiRNAs from both strands are loaded into MEL1. In a 1C/19C - 1G/19G duplex, only the top strand is loaded into MEL1. (d) Comparing two duplexes, 1D/19G - 1C/19H outcompetes 1D/19C - 1G/19H, yielding a 1C/19H MEL1-bound phasiRNA (from the bottom strand).

Fig. S4

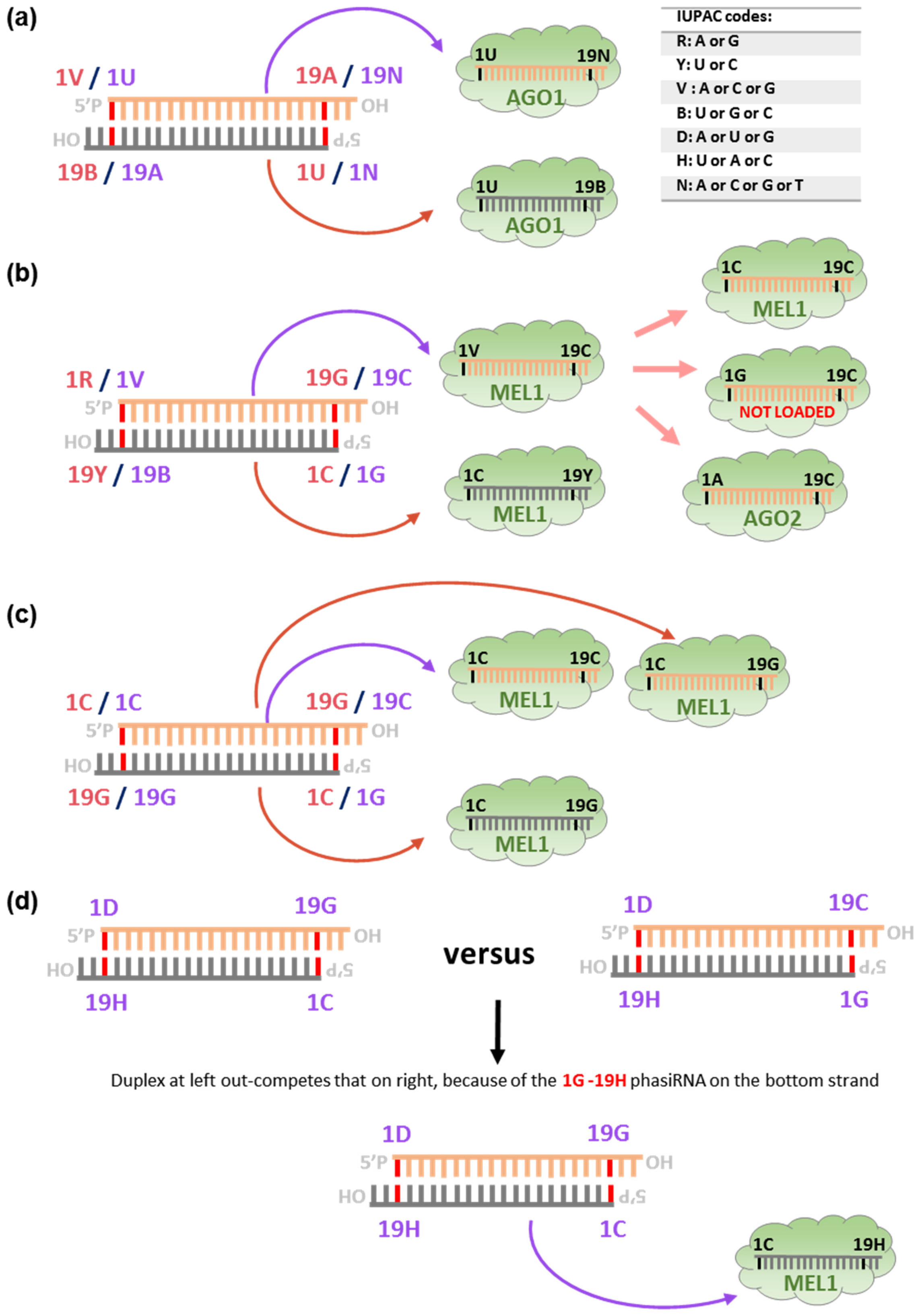

(a)
1V / 1U
19A / 19N
5'P
OH
HO
19B / 19A
1U / 1N
1U
19N
AGO1
1U
19B
AGO1
IUPAC codes:
R: A or G
Y: U or C
V : A or C or G
B: U or G or C
D: A or U or G
H: U or A or C
N: A or C or G or T
(b)
1R / 1V
19G / 19C
19Y / 19B
1C / 1G
1V
19C
MEL1
1C
19Y
MEL1
1C
19C
MEL1
1G
19C
NOT LOADED
1A
19C
AGO2
(c)
1C / 1C
19G / 19C
19G / 19G
1C / 1G
1C
19C
MEL1
1C
19G
MEL1
1C
19G
MEL1
(d)
1D
19G
19H
1C
versus
1D
19C
19H
1G
Duplex at left out-competes that on right, because of the 1G -19H phasiRNA on the bottom strand
1D
19G
19H
1C
1C
19H
MEL1

**Table S1** sRNA libraries from maize and rice used in this study (separate Excel file)

**Table S2** Predicted targets of 21-nt phasiRNAs in rice (separate Excel file)

**Table S3** Predicted targets of 24-nt phasiRNAs in rice (separate Excel file)

**Table S4** Top 30 features, from example comparisons, obtained using information gain.

| **Classification** | **Feature Group** | **No. of features** | **Feature Symbol** |
|---|---|---|---|
| 21-nt phasiRNA vs. P4-siRNA (3' trimmed) | Sequence-based features | 18 | CGG, GG, CG, G, CT, GC, CAT, CGGA, T, ACG, TCGG, AAACG, GGA, C, ACGG, CGGAC, AAC, A |
| | Positional Features | 12 | 1A, 1T, 1C, 19A, 1G, 21G, 21T, 14T, 19C, 20C, 20G, 20A |
| 21-nt phasiRNA vs. miRNA (3' trimmed) | Sequence-based features | 14 | GG, AC, C, G, CT, GCCAA, GTTT, ACTG, GCC, AGA, Shannon Entropy, GC_content, A, GTT |
| | Positional Features | 16 | 1C, 19A, 1G, 2G, 21T, 21G, 19G, 21C, 1T, 19C, 8G, 1A, 8A, 20C, 17T, 14T |
| 24-nt phasiRNA vs. P4-siRNA | Sequence-based features | 23 | CGG, AT, GC_content, ACGG, CG, GG, G, GAC, CGGA, GGAC, T, GGA, GGC, CAT, ACG, GACG, TTG, TAT, TCA, CGGAC, AC, TT, GA |
| | Positional Features | 7 | 1A, 1T, 22C, 23T, 22T, 22A, 1C |

Note: Sequence-based features denote the frequency of the k-mer motifs as they appear along the length of the sequence. Positional Features denote the presence or absence of a given nucleotide at that position in a sequence.

**Method S1** Dataset used for cross validation study

We used several representative positive and negative datasets to perform a five-fold cross validation. The use of these sets in the cross-validation steps allowed us to compare the performance of the classifiers with these different datasets. The positive sets consisted of non-redundant sequences of reproductive phasiRNAs, including sets of phasiRNAs whose length is either 21- or 24-nt. The sets characterized by either sequence length comprise phasiRNAs from previously-identified *PHAS* loci published for rice and maize. Specifically for rice, we utilized small RNA libraries published by Fan et al., 2016 and analyzed these libraries using our previously published python-based FASTQ processing script to produce the 'tag count' formatted files (Patel *et al.*, 2015). Next, we identified 2024 21-nt *PHAS* loci and 59 24-nt *PHAS* loci from rice using a tool called *PHASIS* (Kakrana *et al.*, 2017), and we used 463 21-nt *PHAS* loci and 176 24-nt *PHAS* loci previously described for maize (Zhai *et al.*, 2015) to obtain 21-and 24-nt phasiRNAs. From these loci, we identified non-redundant sets of 20240 (rice) and 9260 (maize) premeiotic 21-nt phasiRNAs, and 2950 (rice) and 8800 (maize) meiotic 24-nt phasiRNAs; these comprise the small RNAs from these *PHAS* loci that were represented in the sequencing data from the reproductive tissues. To ensure high quality of the data, we used in our cross validation studies for training/testing the classifiers only the top 1000 most-abundant phasiRNA sequences from each species represented within the sets of sequences of either length. The negative training sets were built by gathering several types of non-redundant sRNA sequences. For simplicity, we refer to all of them as *non-phasiRNAs*, as each sequence of each type was selected based on the criterion of not containing reproductive phasiRNAs

- Mature miRNA sequences were downloaded from miRBase (Kozomara & Griffiths-Jones, 2014), version 21. We combined miRNAs from rice (n=553) and maize (n=203), totaling 756 miRNAs. We trimmed miRNAs longer than 21-nt to 21-nt, trimming from the 3'- end to obviate any length differences in comparison to the 21-nt phasiRNAs. Thus, we used this set (n=756) only while classifying 21-nt phasiRNAs.

- P4-siRNAs sequences were selected from either rice or maize small RNA libraries (Table S1). These were selected using a length filter of 24 nucleotides, and the criterion that the sequence should match the genome more than 10 times and overlaps with repetitive regions, consistent with repetitive origins. As an additional filter to avoid reproductive phasiRNAs, the P4-siRNAs were from vegetative tissues in which reproductive phasiRNAs are absent (Johnson *et al.*, 2009). In the libraries we analyzed, using these criteria, we identified 14008 non-redundant P4-siRNAs from maize and 12530 from rice as our data set. Of these, we selected the top 1000 most abundant sequences from rice and separately from maize. As with the miRNAs, for comparisons to 21-nt phasiRNAs, we trimmed P4-siRNAs to 21-nt from the 3'- end. For comparisons to 24-nt phasiRNAs, no trimming was required since the P4-siRNAs were preselected as 24-nt small RNAs.
- Small RNAs from tRNA and rRNA loci in either rice or maize were extracted from the vegetative libraries described above. The tRNA loci in both genomes were identified using tRNAscan-SE (Lowe & Eddy, 1997) with default parameters. The rRNA loci were identified using RepeatMasker (Smit *et al.*, 2013) with a default cut-off score of 225. The small RNAs derived from these loci were combined for the two genomes (since tRNAs and rRNAs are highly conserved between rice and maize), and we randomly selected 500 tRNAs and 500 rRNAs.

Finally, to assess the predictive performance of the algorithm on new sequences, we selected the remaining 8260 and 19240 21-nt phasiRNAs and 7800 and 1950 24-nt phasiRNAs from maize and rice, distinct from those that comprised the positive set used for cross validation study. In addition, we also generated 18 sRNA libraries from *Setaria viridis*, a grass, in order to use reproductive phasiRNAs from *S. viridis* to test predictive sensitivity across species.

We identified 1593 21-nt *PHAS* loci and 381 24-nt *PHAS* loci from *S. viridis* using *PHASIS* (Kakrana *et al.*, 2017). From these loci, for subsequence cross-species comparisons, we used non-redundant sets of 2000 premeiotic 21-nt phasiRNAs, and 2000 meiotic 24-nt phasiRNAs.

**Method S2** Features included in the machine learning algorithm, and their selection

The machine learning classification pipeline integrated both sequence-based and structural features, as follows:

**Sequence Based Features:**

1. The frequency of k-mer motifs: We computed normalized frequencies of all 1 to 5 nt k-mer patterns (see patterns below) using a sliding-window of length k (k = 1, 2, 3, 4, 5). A particular string of length k could slide along the length of the sequence, denoted as L, by a step of 1 nt. If the string in the sequence matched with some pattern *i* within the window, the count of that pattern in the sequence, denoted as $C_i$, was increased by one, using the following formula to calculate frequencies:

   $F_i = C_i / S_k$, for i = 1 to 1364 , where $S_k = L - k+1$, and $S_k$ is the total number of times the sliding-window of length k could slide along the sequence.

   The frequencies of all 1 to 5 nt k-mer patterns (4+16+64+256+1024=1364 patterns) were as follows:

    1. 4 one-mer patterns (A,C,G,U):
    2. 16 two-mer patterns (AA,AC,AG,AU,…,UU)

3. 64 three-mer patterns (AAA,AAC,AAG,AAU,…,UUU)
4. 256 four-mer patterns (AAAA,…,UUUU)
5. 1024 five-mer patterns (AAAAA,…,UUUUU)

Thus, phasiRNAs and non-phasiRNAs were characterized as a vector consisting normalized frequencies of 1364 k-mer motifs. We used the repDNA package (Liu *et al.*, 2015) in Python to generate these features.

2. GC_content = (|C|+|G|) / L*100, where |C| and |G| denote the number of nucleotides C or G in the sequence, respectively.

**Positional Features:**

We computed a position-specific base usage vector by simply representing a sequence in a numerical fashion, denoting a presence (1) or absence (0) of each nucleotide at each position in the whole sequence. Each position in the sequence can have four possibilities for each of the four nucleotides (A, C, G, and T). A sequence of length L will have a position feature vector of length 4*L, where 4 is the number of nucleotides and L is the length of a sequence. For example, a sequence of length 21 will be a vector of 4*21=84, such as the following:

Positional feature vector = < 1A,1C,1G,1T,2A,2C,2G,2T,…,21A,21C,21G,21T >

Each of the entries in the above vector holds a value of either 1 (presence of a given nucleotide at that position) or 0 (absence of a given nucleotide at that position). The length of sRNA sequences in our dataset is not constant. Therefore, we used a length of the longest sRNA found in the dataset as an upper limit to generate the positional features. When sequences of identical lengths were compared, for example 24-nt phasiRNAs versus 24-nt P4-siRNAs, no length adjustment was required. In other cases, classification models were built from sequences of mixed lengths; for example, in some of our data sets, the longest read was from a tRNA, and 33 nt in length. Therefore, for each sequence in those models, we obtained a positional vector of length 4*33=132. For sRNA sequences shorter than 33 nt, the extra positions in the positional feature vector are all 0.

**Shannon entropy:**

Shannon entropy score is an indicative measure of information content, assessing the repetitiveness in a sequence. We computed the Shannon entropy for given symbol frequencies for given sequence. The formula for calculating Shannon entropy score is as follows:

$$S(L) = - \sum f(x) \log2 (f(x)),$$

where L is a sequence length and f(x) is the frequency of the symbol (x - A, C, G, or T) in the sequence. The lower the value of entropy, the higher the repetitiveness is in a sequence. We used a length-normalized Shannon entropy score calculated as follows:

$$\text{Normalized Shannon entropy score} = S(L) / L.$$

Thus, for each sequence in the positive and the negative sets, we calculated 1498 features (1365 sequence-based + 132 positional-based + Shannon entropy).

To eliminate redundancy from among the 1498 features, the attribute selection tool of WEKA was used prior to performing classification, separately for each positive vs negative set comparison. Information gain (also known as entropy) was employed to evaluate the value of each feature, for either the phasiRNA or non-phasiRNA classes. Values of information gain (i.e. the relative contribution) for each feature varied from zero (no information) to one (maximum information); features conveying more information thus had a higher information gain value. Features with lower values could be removed, with the aim to reduce the dimensionality of the feature set by using a minimum number of maximally informative features. Thus the features were ranked by information gain, and the top 250 were selected (Fig. S1 a,b), as this subset included 95 to 97% of informative features (Fig. S1e), reducing dimensionality without compromising or negatively impacting the classifier's performance. As examples of our feature selection process, we listed the top 30 out of 250 selected features in Table S4.

**Method S3** Generation and computational analysis of sequencing data from maize and *Setaria viridis.*

Four biological replicate libraries were constructed from the *rdr2/mop1* maize mutant (Alleman *et al.*, 2006), using young ears (3 to 5 cm in length); the materials were generously donated by Drs. Mario Arteaga-Vazquez and Vicki Chandler (formerly of the University of Arizona). Small RNA library construction was as previously described (Lu *et al.*, 2007). A total of 18 libraries from *Setaria viridis* were constructed using the Illumina TruSeq sample preparation method. We used two biological replicate libraries from the leaf, whole panicles (inside leaf sheath), whole panicles (coming out of leaf sheath), whole panicles (completely out of leaf sheath), whole panicles (completely out of leaf sheath, after pollination), spikelet (inside leaf sheath), spikelet (coming out of leaf sheath), and spikelet (completely out of leaf sheath). Small RNA libraries were processed using a custom python-based FASTQ pipeline (Patel *et al.*, 2015). Briefly, the processing script removes adaptors and low-quality reads, and retains sequences with lengths between 18 and 31 nt for the maize data and 18 and 34 nt for *S. viridis* (the difference is because the maize data were generated using shorter total sequence reads). Then, these sequences were mapped to the respective genome: maize version AGPv2 or *S. viridis* version v1 (from JGI).

Mapping was performed using Bowtie (Langmead *et al.*, 2009). Reads that perfectly matched (no mismatch) to the maize genome and *S. viridis* genome, excluding those matching to structural RNAs (tRNAs or rRNAs) were used for further study.

**Method S4** Extraction of a set of maize 22-nt hc-siRNAs

The 22-nt, putative heterochromatic siRNAs that are RDR2-independent, and thus far found only in maize (Nobuta *et al.*, 2008) were selected from the maize small RNA libraries (Table S1). These were selected using a length filter of 22 nucleotides, and the criterion that the sequence should overlap with highly repetitive regions (i.e. hits > 50 in the genome), consistent with repetitive origins.
